# The impact of ancestral, environmental and genetic influences on germline *de novo* mutation rates and spectra

**DOI:** 10.1101/2024.05.17.594464

**Authors:** O. Isaac Garcia-Salinas, Seongwon Hwang, Qin Qin Huang, Joanna Kaplanis, Matthew D.C. Neville, Rashesh Sanghvi, Felix Day, Raheleh Rahbari, Aylwyn Scally, Hilary C. Martin

**Affiliations:** Wellcome Sanger Institute, Wellcome Genome Campus, Hinxton, CB10 1SA, UK; Department of Genetics, University of Cambridge, Cambridge, UK; MRC Biostatistics Unit, School of Clinical Medicine, University of Cambridge, Cambridge CB2 0SR, UK; Genomics England, London, UK; MRC Epidemiology Unit, Box 285 Institute of Metabolic Science, University of Cambridge School of Clinical Medicine, Cambridge CB2 0QQ, UK

**Author notes:** Joint first authors (O. Isaac Garcia-Salinas, Seongwon Hwang). Joint senior authors (Raheleh Rahbari, Aylwyn Scally, Hilary C. Martin).

## Abstract

D*e novo* germline mutation is an important factor in the evolution of allelic diversity and disease predisposition in a population. Here, we study the influence of genetically-inferred ancestry and environmental factors on *de novo* mutation rates and spectra. Using a genetically diverse sample of ∼10K whole-genome sequenced trios, one of the largest *de novo* mutation catalogues to date, we found that genetically-inferred ancestry is associated with modest but significant changes in both germline mutation rate and spectra across continental populations. These effects may be due to genetic or environmental factors correlated with ancestry. We find epidemiological evidence that exposure to tobacco smoke is significantly associated with increased *de novo* mutation rate, but it does not mediate the observed ancestry effects. Investigation of several other potential mutagenic factors using Mendelian randomisation showed no consistent effects, except for age of menopause, where increased age corresponded to a reduction in *de novo* mutation rate. Overall, our study presents evidence on new factors influencing *de novo* mutational rate and spectra.

## Main text

Germline mutation is a fundamental evolutionary process, and *de novo* germline mutations are a major cause of developmental disorders ^1^. Such mutations occur at an exceptionally low rate ^2,3^, but this rate can be influenced by exposure to mutagens such as chemotherapeutic agents ^4^ and ionising radiation ^5^. Tobacco smoke is known to affect the accumulation of *de novo* micro/minisatellites ^6^, but its effects on *de novo* point mutations, the best-characterised form of genetic variation, have not yet been studied. Genetic factors may also influence germline mutation rate ^7^. Rare variants in DNA repair genes are well-known modifiers of somatic mutation rates and spectra ^8–10^, and have been shown to contribute to elevated rates of germline mutation ^4,10^. Certain common variants are associated with the rates of germline mutations at microsatellites ^11^, and common variants that decrease age of menopause, many of which implicate DNA repair genes, are associated with increased rates of *de novo* point mutations in the female germline ^12^. Human polymorphism data indicate that mutation spectra have varied over time and between human populations ^13,14^, and a variety of genetic and environmental causes have been proposed for this ^15^. However, the study of ancestral, environmental, and genetic factors influencing directly-measured germline mutations has been constrained due to the lack of appropriate datasets. In this paper we explore the influence of continental-level genetic ancestry, smoking behaviour, and parental genetics on the *de novo* point mutation rate by analysing over 10,000 whole-genome sequenced parent-offspring trios from the Genomics England 100,000 Genomes project.

We used *de novo* single nucleotide variants (henceforth referred to as “DNMs”) from 10,557 family trios which had been previously called and filtered as described elsewhere ^4^. These data consisted of 742,753 high confidence DNMs (average 70.35 per trio). Around 26% of these DNMs (n = 197,567; average 19.51 per trio) had been phased to their parent-of-origin using a read pair-based approach ^4^.

We first explored whether the mutation rate and spectra differed between individuals of different genetic ancestries, using the continental-level genetic ancestry classifications produced by the 100,000 Genomes project ^16^. These classifications are based on genetic similarity to individuals of known origins from the 1,000 Genomes Project ^17^. Although this classification does not fully capture the genetic diversity of human populations, it suffices for the main purpose of our analyses. For this analysis, we included 9,831 trios for which both parents were inferred to come from the same continental-level ancestry group (198 African [AFR], 216 American [AMR], 53 East Asian [EAS], 8,102 European [EUR], and 1,249 South Asian [SAS]). First, we tested the association between ancestry and total DNM rate using generalised linear models, controlling for parental ages and technical covariates associated with the ability to call DNMs (**<u>Methods</u>**). We detected significant DNM rate differences (false discovery rate, FDR<5%) between the AFR and EUR groups (rate ratio=1.05, p=3.51e^-6^), AFR and AMR groups (rate ratio=1.037, p=1.16e^-2^), AFR and SAS groups (rate ratio=1.035, p=1.78e^-3^), and EUR and SAS groups (rate ratio=0.98, p=3.02e^-3^) (**Figure 1A; <u>Supplementary Table 1</u>**).

**Figure 1.**
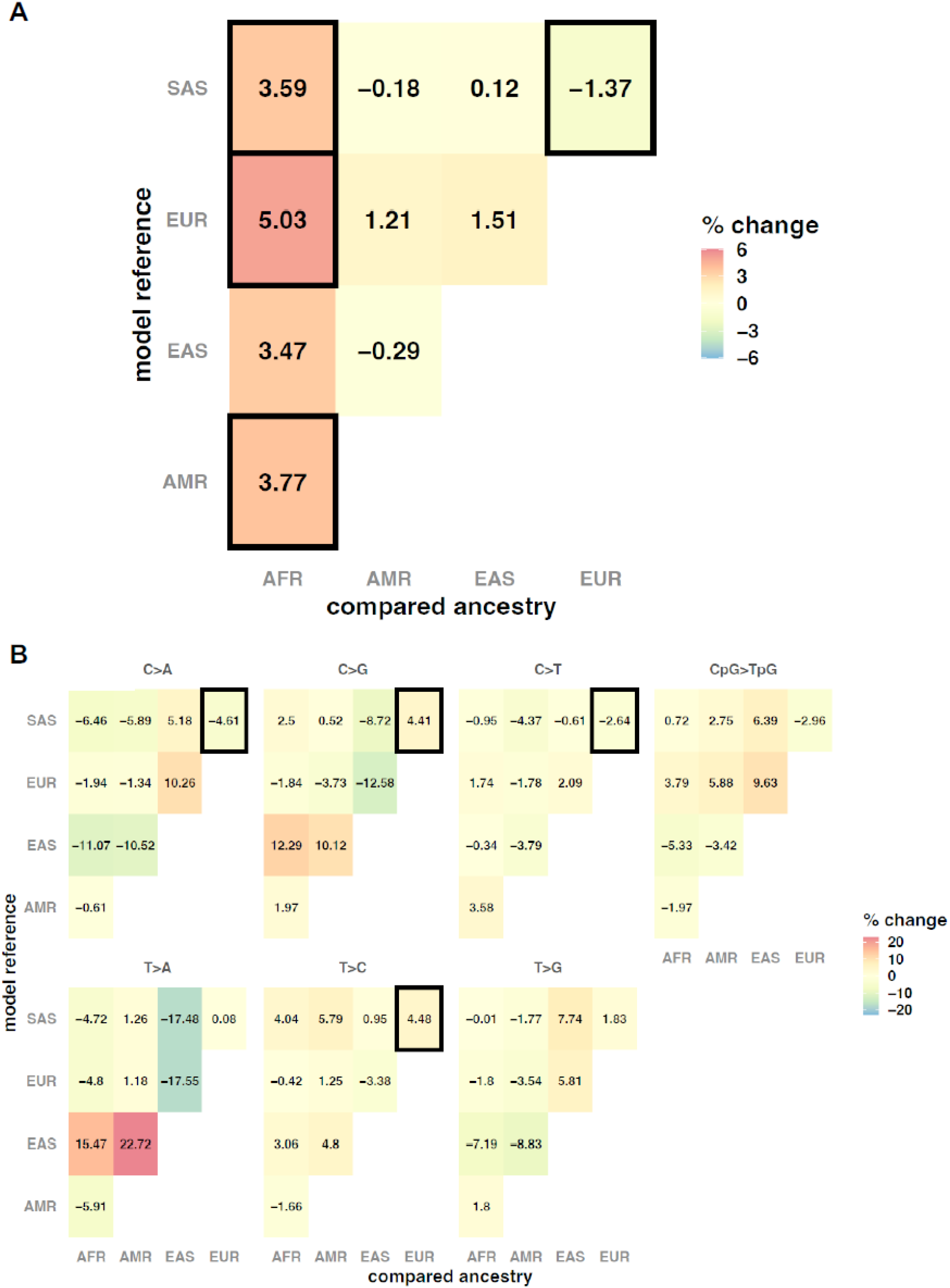
Pairwise ancestry comparisons for a) DNM rate and b) DNM spectra (pyrimidine substitution proportions). Heatmap colours and values correspond to the percentage change in DNM counts (or pyrimidine proportions) expected for the ancestry group indicated on the X-axis when comparing to the model reference indicated on the Y-axis. Percentage changes are obtained from the rate ratio effect estimate from each regression model. Decimals in percentage values have been rounded to a double-digit precision for plotting purposes. Bold squares indicate effect significance at 5% FDR (p adjusted <= 0.05).

As allele frequencies were not among the criteria to call or filter DNMs ^4^, these ancestry differences in DNM rate could not be driven by differences in the numbers of individuals from different ancestry groups in reference databases. However, in theory, these observed differences in DNM counts could be confounded by average differences in read mapping quality between ancestry groups due to reference bias ^18^. We controlled for several metrics to test if mapping and other technical biases may be affecting our observed associations with ancestry, but found no evidence of this (**<u>Supplementary Note 2</u>; Supplementary Figures 1 and 2)**. Similarly, a sensitivity analysis controlling for the number of protein-altering DNMs suggested that our findings were not the result of particular genetic ancestries being enriched for pathogenic DNMs, which could in theory happen if there were ancestry-correlated recruitment biases into the 100,000 Genomes Project (**<u>Supplementary Note 2</u>, Supplementary Figure 2**).

We then tested whether ancestry was associated with differences in mutational spectra. For this, we divided the DNMs into seven categories according to which pyrimidine substitution was involved (and whether it was at a CpG site in the case of C>T transitions^19^), and then calculated the proportion of mutations in each category per trio (**<u>Methods</u>**). We ran generalised linear regression models to test for pairwise differences in these substitution proportions between ancestry groups. We identified significant differences (FDR<5%) in the proportion of several types of mutations between the EUR and SAS groups, namely for C>A (rate ratio = 0.95, p = 3.015e^-4^), C>G (rate ratio = 1.044, p = 2.07e^-3^), C>T (rate ratio = 0.97, p = 2.01e^-3^), and T>C (rate ratio = 1.044, p = 8.82e^-7^) mutations (**Figure 1B, <u>Supplementary Table 2</u>**). The differences in C>A and T>C proportions recapitulate what has been previously reported ^14^ (**<u>Supplementary Note 3</u>**).

These ancestry differences in DNM rates and spectra could reflect genetic differences between ancestry groups, or differences in environmental exposures between them, which could include, for example, differences in diet or exposure to common mutagens such as cigarette smoke ^20^. Very limited data on environmental exposures were available in the 100,000 Genomes Project, but we assessed the effect of smoking on DNM rate using the electronic health record (EHR) data (**<u>Methods</u>**). Using ICD10 codes, we created a binary “*ever smoked*” phenotype per individual. We then re-analysed associations between total DNM count and ancestry in a set of non-admixed trios in which at least one parent had ICD10 data available (192 AFR; 203 AMR; 46 EAS; 7,387 EUR; 1,203 SAS). For this, parental smoking was classified as: both parents smoke (n = 293), only the father smokes (n = 664), only the mother smokes (n = 833), or neither parent smokes (n = 7,241). We obtained nearly identical ancestry effects to those reported in **Figure 1** (**Supplementary Figure 3**), indicating that differences in parental smoking behaviour across ancestries are unlikely to be driving these ancestry associations. We observed significant effects of having one or both parents smoke on total DNM count (**Supplementary Figure 4**), with the caveat that these effect sizes may be noisy, since in many trios the smoking status of one parent was unknown and they were assumed to be a non-smoker. To refine our estimates of the smoking effect, we restricted to a set of 6,599 fathers and 9,133 mothers with the relevant subset of EHR data, and tested the effect of smoking on the number of DNMs derived from the relevant parent (i.e. phased DNM count, **Methods**). We found that having ever smoked was a significant predictor of increased DNM rate in females (rate ratio = 1.038, p = 2.5e^-2^) and males (rate ratio = 1.019, p = 4.27e^-2^), and in a sex-combined analysis (rate ratio = 1.024, p = 3.6e^-3^, **Figure 2**).

**Figure 2.**
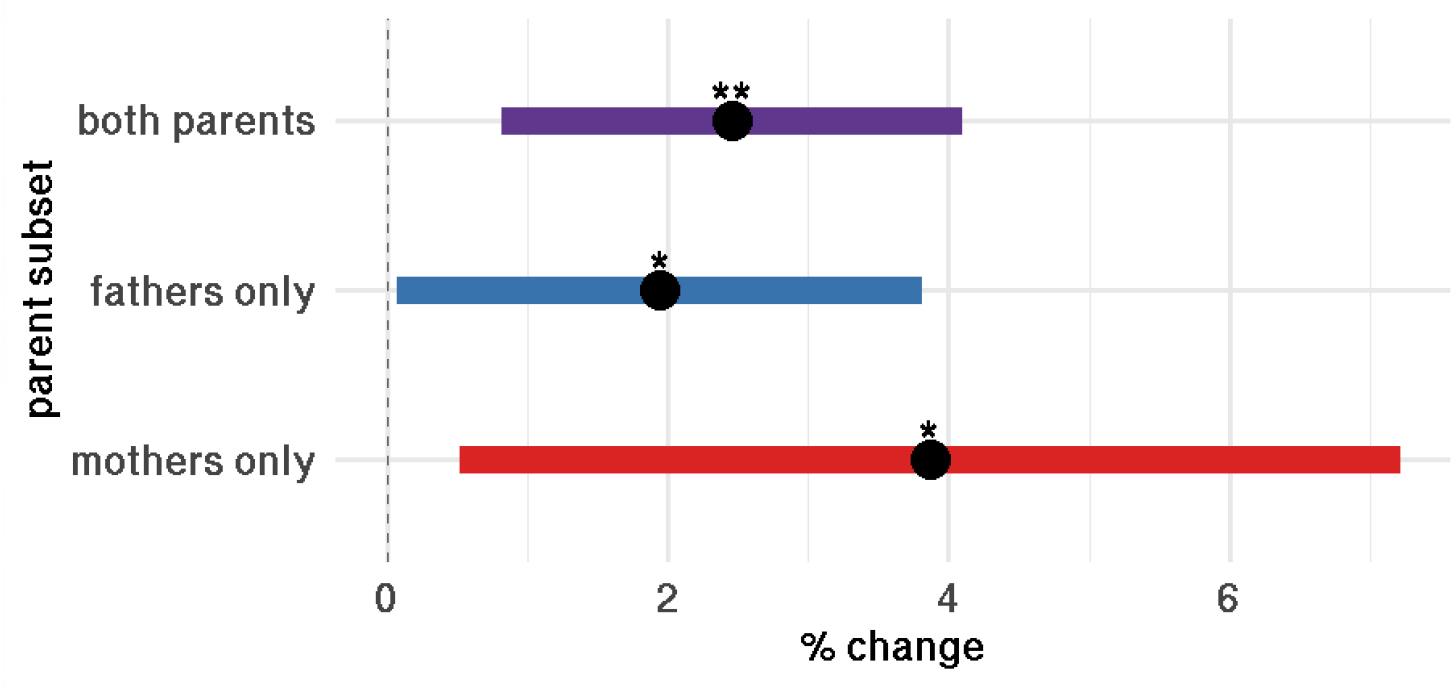
Effect of smoking on DNM rate, using phased DNMs from 6,599 fathers and 9,133 mothers with EHR data. The phenotype was derived from Z58.7 and F17 codes ICD10 codes corresponding to “*exposure to tobacco smoke*”, and “*mental and behavioural disorders due to use of tobacco”*, respectively. The y-axis indicates whether mothers, fathers or both were used in the analysis. Percentage changes are obtained from the rate ratio effect estimate when comparing “ever smoked(1)” vs “ever smoked(0)” in the regression model. Error bars show 95% confidence intervals. *p<=0.05, **p<=0.005.

Finally, we attempted to identify differences in mutation spectra associated with parental smoking behaviour by applying different methods and definitions of mutation spectra (**<u>Supplementary Note 4</u>**). We did not identify any significant associations between smoking and specific substitutions types, and neither were we able to identify any known mutational signature ^21^ associated with smoking on DNMs (**<u>Supplementary Note 4</u>, Supplementary Figure 5**).

Within the subset of unrelated parents inferred to have European genetic ancestry (7,786 mothers, 7,692 fathers), we estimated the variance in DNM rate explained by variants with minor allele frequency >=0.1%, using GREML-LDMS ^22^. For this we used the parentally phased DNM counts previously produced by Kaplanis et al.^4^. After accounting for parental age and technical factors (**<u>Methods</u>**), we did not obtain a significant SNP-heritability estimate with any of the minor allele frequency-linkage disequilibrium (MAF-LD) bin cutoffs tested, in either fathers, mothers, or both combined (**Figure 3**). From this, we concluded that variance explained by common variants on DNM rate must be too low to be detected in this sub-cohort, given its sample size. However, we note that potential effects of ancestry-associated genetic variation on DNM rate would be excluded from this analysis.

**Figure 3.**
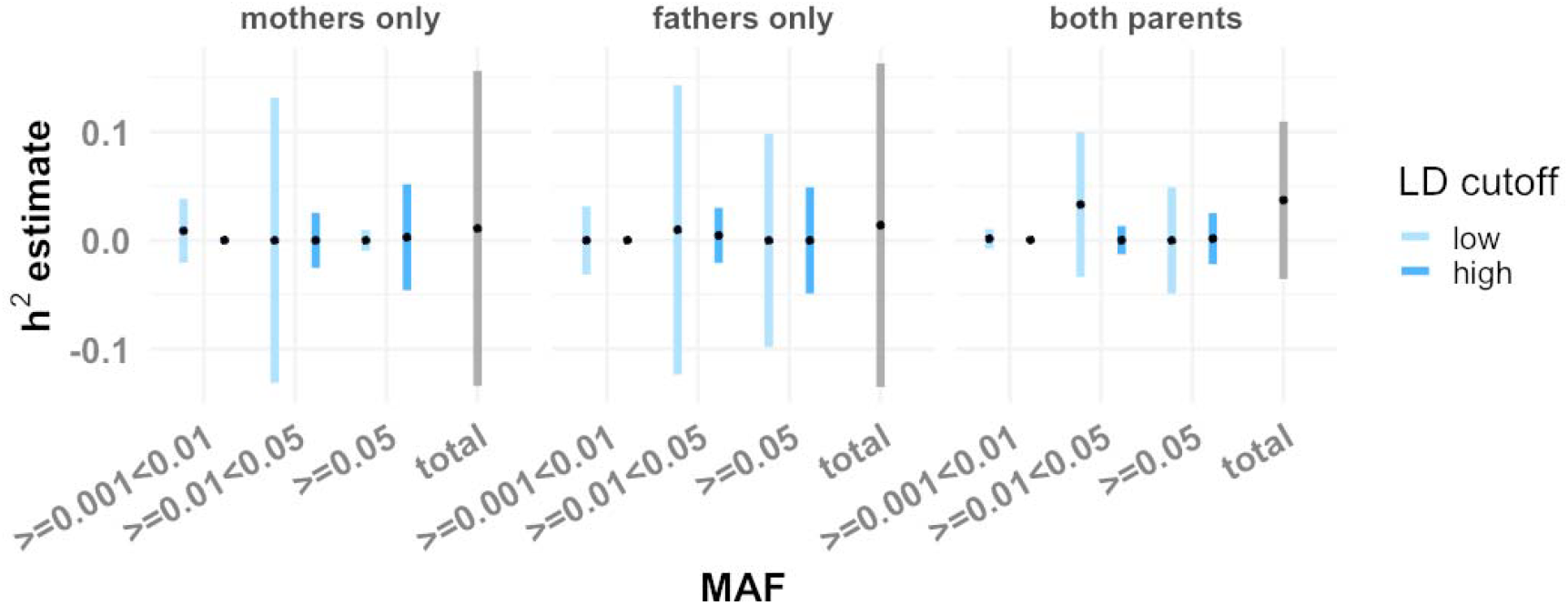
Estimates of SNP heritability of maternally- and paternally-derived DNM counts, from GREML-LDMS. Left plot shows heritability estimates of maternally phased DNMs in the mothers subset alone. Centre plot shows estimates for paternally phased DNMs in the father subset alone. Right plot shows estimates from both parents combined. Error bars show 95% confidence intervals. Estimates were divided by MAF and according to whether the LD score of the variant was above or below the genome-wide LD median.

We also ran GWAS on a broader sample including related European-ancestry individuals (7,993 mothers, 7,892 fathers) while controlling for relatedness using SAIGE ^23^. This did not detect any genome-wide significant SNPs or indels (p <= 5×10^−8^) in any sample subset (**Supplementary Figure 6**).

We applied two-sample Mendelian Randomisation (MR) analyses to explore the influence of a range of factors on DNM rate, including some associated with reproductive traits. The exposures considered included age at natural menopause (ANM) (which we took as a positive control ^12^), three smoking-related measures ^24^, alcohol use ^24^, body mass index (BMI) ^24^, and three traits chosen based on their associations with the top SNP from our DNM GWAS, namely sleep duration, hydrocele and spermatocele, and diseases of male genital organs (see **<u>Methods</u>**). Following standard MR procedures ^25^, for each exposure of interest, we selected SNPs as instrumental variables based on a p-value threshold 5×10^−8^ from publicly available GWAS, and applied LD clumping (r^2^>0.1) to identify independent SNPs (**<u>Supplementary Table 3</u>**). ANM was found to have a negative causal effect on DNM rate in mothers but not fathers, as previously reported in the same dataset ^12^ (**Supplementary Figure 7**). Importantly, its effects on maternal DNM rate were consistent across MR methods. We did not detect any further significant causal effect estimate for any of the remaining exposure traits, except that the number of drinks per week had a nominally significant causal effect on lower DNM count in fathers when using the simple median MR method (**Supplementary Figure 7**). However, this result was not significant in any of the three other MR methods, and so should be treated with caution.

To conclude, in this work we explored the association of genetic and environmental factors with the genome-wide rate of point mutations in the germline. Although it is suspected that genetic factors play a role influencing the DNM rate ^26,27^, we failed to detect any significant SNP heritability in this cohort attributable to variants down to a frequency of 0.1%, consistent with a previous smaller study ^28^. The observation that common SNPs associated with age of menopause are causally associated with DNM rate in mothers ^12^ (**Supplementary Figure 7**) implies that there are common variants influencing DNM rate, but that they likely explain very little phenotypic variance. The error bars on our estimates in **Figure 3** imply that the heritability attributable to common SNPs with frequency greater than 5% is likely to be less than 15%. One possible explanation for the small contribution of common variants is the strong selection pressure against large-effect mutator alleles in the germline progenitors which prevents them from acquiring potentially deleterious new variants ^29^. Future studies with larger sample sizes should investigate whether rare variants genome-wide explain an appreciable fraction of the variance in DNM rate.

We have presented direct evidence that smoking is associated with the number of *de novo* point mutations in humans (**Figure 2**). Specifically, we find that being a smoker increases the DNM count by ∼2%, equivalent to less than one extra DNM per smoking parent over the reproductive lifespan. We emphasise that our effect size estimates should be treated with caution since they rely on smoking behaviour being reported accurately in the health records, and it is plausible that these records are biased towards recording heavy rather than occasional smoking behaviour. We also note that these associations do not show that smoking is *causal* for increased DNM rate, and indeed, both the MR and signature analyses failed to provide evidence of this (**Supplementary Figure 7**), although they may simply be underpowered; conceivably, smoking could be correlated with exposure to other mutagens.

We also show direct evidence that the present-day DNM rate differs between continental ancestry groups (**Figure 1A**), being ∼3-5% higher in AFR than EUR, SAS and AMR, and ∼1% lower in EUR than SAS. Although we show that these differences are unlikely to be due to differences in smoking behaviour, they may be due to differences in other environmental exposures and/or ancestry-associated genetic variation. The finding of a difference in DNM rate between EUR and SAS is supported by significant differences in mutational spectra between these two groups (**Figure 1B**). Compared to a previous study of variation in mutational spectra across continental populations ^14^, we found a similar C>A depletion and T>C enrichment, when contrasting EUR with SAS ancestry. However, unlike that study, we also found C>G and T>C enrichment in EUR, but no significant enrichment for T>G (**<u>Supplementary Note 3</u>**). Also despite significant differences in overall DNM rate between AFR and other groups, we found no significant differences in mutational spectra, whereas many such differences were found by Harris and Pritchard, 2017. These discrepancies may be because the polymorphism data they analysed reflect mutations that occurred over thousands of generations rather than just one, and the shifts in mutational spectra may stem from transient differences in environmental or genetic causes of DNMs that no longer exist between contemporary human ancestry groups. We note that the small average differences in DNM rate between ancestry groups that we have reported are likely to be exceeded by differences due to variation in the distribution of parental age at birth ^30^, which is changing across time ^31^ and which has a much larger influence on DNM rate. Indeed, branch-length comparison between African and non-African populations has previously suggested an increase in mutation rate in non-Africans since out-of-Africa ^32^. Future studies may need to consider whether conclusions are changed by integrating these or other ancestry-associated differences in mutation rate and spectra into population genetic models ^33^, inferences about ancestry and demographic history (e.g. ^34^), and mutation rate models used for inferring genic constraint ^35^ and discovering genes enriched for DNMs in disease cohorts ^1,36^.

Finally, this work demonstrates the imperative to include greater diversity of genetic ancestry in studies of DNMs, and to collect comprehensive epidemiological data on potential environmental mutagens. This will enable more powerful and robust investigation of the genetic and environmental influences on germline mutation. Since existing methods for detecting and estimating genetic associations rely on genetically homogeneous cohorts, future studies may also require methodological developments to effectively disentangle ancestry-correlated genetic variants from environmental factors affecting DNM rates.

## Acknowledgements

This research was made possible through access to data in the National Genomic Research Library, which is managed by Genomics England Limited (a wholly owned company of the Department of Health and Social Care). The National Genomic Research Library holds data provided by patients and collected by the NHS as part of their care and data collected as part of their participation in research. The National Genomic Research Library is funded by the National Institute for Health Research and NHS England. The Wellcome Trust, Cancer Research UK and the Medical Research Council have also funded research infrastructure. We also thank Daniel Malawsky and Federico Abascal for their valuable feedback and time for discussion. This research was funded in part by Wellcome (grant no. 220540/Z/20/A, “Wellcome Sanger Institute Quinquennial Review 2021–2026”). For the purpose of open access, the authors have applied a CC-BY public copyright licence to any author accepted manuscript version arising from this submission.

## Author Contributions

OIGS and SH conducted the analyses and drafted the manuscript. QQH, JK, MDCN, RS and FD advised on QC or specific analyses. RR, AS and HCM supervised the study.

## Materials and methods

### Whole genome *de novo* mutations and phasing information across a cohort of 13,949 family-trios

We used data from the rare disease branch from the 100,000 Genomes Project (100kGP) from Genomics England ^37,38^. Studied families were selected on the basis of having at least one offspring with an undiagnosed rare disease ^37,38^. Although 13,949 trios were originally recruited as part of this, the data freeze we used (v16) contained information for just 12,017 trios due to some individuals having withdrawn their consent to participate. *De novo* variant calls (DNMs) per trio were generated using the Platypus variant caller ^39^ and included single nucleotide variants (SNVs, n = 906,643) and short insertion-deletion events of up to 250bp (INDELs, n = 72,481), which were previously filtered using a stringent criteria that is fully described elsewhere ^4^. *De novo* SNVs were phased to the parent of origin using parental heterozygous single nucleotide polymorphisms found in a 500bp vicinity of the reported DNM. This strategy allowed phasing of 26% of the original DNMs, out of which 80% phased to the paternal germline ^4^. Downstream analyses were focused on *de novo* SNVs, and we refer to these as DNMs for simplicity.

### Genetically inferred ancestry information

Ancestry information for all of the participants was readily available in the Genomics England (GEL) research environment. This resource was produced from joint principal component analysis (PCA) from the 100kGP individuals and the 1000 Genomes Project phase 3 (1kGP3) ^37^. Briefly, PCs for the 1kGP3 were calculated using a set of 188,382 high quality SNPs (MAF >= 0.05) intersecting with the 100kGP, in unrelated individuals. The 100kGP data was then projected onto 1kGP3 PC loadings, and a random forest classifier based on the first 8 PCs and continental-level ancestry classifications from 1,000 Genomes was used to predict ancestry for each individual in the 100kGP dataset. Ancestry classifications from 1,000 Genomes corresponded to one of the five continental-level super-populations: African (AFR), American (AMR), European (EUR), East Asian (EAS), and South Asian (SAS) ^17,37^. Each individual in the 100kGP cohort was assigned a probability of belonging to each of these populations.

Finally, unrelated individuals assigned to each population with a probability >= 0.8 were used to calculate population-specific PCs ^37^. European-specific PCs were used in the heritability estimation and GWAS, which were focused on individuals inferred to have European ancestry.

### Sample filtering

We removed nine trios in which the proband was previously identified as having a significantly elevated rate of DNMs (hypermutator individuals)^4^, to prevent spurious associations in our study due to their elevated DNM rate and characteristic DNM spectra. Some families in this study included multiple offspring. As the DNM rate increases linearly with age ^2,3^, we kept the trio corresponding to the youngest sibling in these multiplex families to increase our chance of observing DNM events. We further removed all trios with >= 1 individuals missing metadata on date of birth, sequencing statistics, or *de novo* mutation calling Bayes factors. This left 10,557 trios for analysis.

### Associations between ancestry and DNM rate

For this analysis, DNM rate was defined as the total DNMs detected per trio (i.e. unphased DNMs). For ancestry, we relied on the classifications readily available in the GEL research environment (described above). For each individual, we took the ancestry assignment with the highest probability (see previous section). We further filtered out trios with parents assigned to different ancestry groups, leaving 9,833 non-admixed trios for analysis. DNM counts are influenced by parental sex and parental age at conception ^2^, as well as by technical factors influencing the ability to call DNMs (fully described in Kaplanis et al., 2022), so our models account for these.

Associations were run using generalised linear models (R glm() function) from the quasi Poisson family (logit link), (selected to account for the mean-variance overdispersion of the phased DNM count data, **<u>Supplementary Note 1</u>**), as shown in **Model 1**:

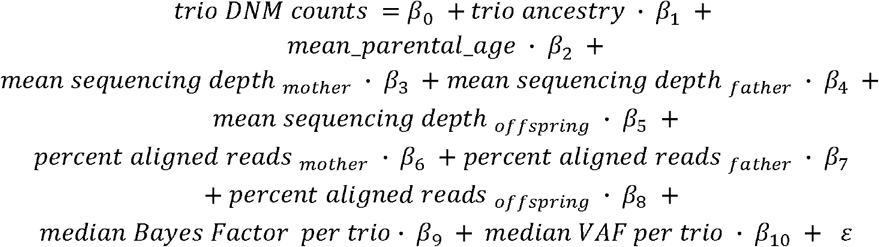

Here, “*trio ancestry*” is a factor covariate taking one of 5 possible identities (AFR, AMR, EAS, EUR, SAS), and “*median VAF per trio*” corresponds to the median offspring variant allele fraction of all high quality DNMs found in such an offspring. As ancestry is a five-level factor covariate, a single ancestry was used as a baseline to compare against all of the remaining four. We iteratively changed this baseline until we obtained all possible, non-redundant, pairwise ancestry comparisons (n = 10). We took the exponential of each ancestry coefficient to obtain rate ratios (RRs). In this context, RRs represent the change associated to a given ancestry against the baseline. All p-values associated with ancestry coefficients were corrected to account for multi-testing using the R p.adjust() function and the FDR method.

### Associations between ancestry and DNM spectra

The mutational spectrum was defined as follows. Each DNM was classified according to the pyrimidine base of the Watson-Crick base pair, which allows for a standardised way to identify and compare mutational patterns ^40^. In this way, each mutation can be classified as one of six possible pyrimidine substitutions (i.e. C>A, C>G, C>T, T>A, T>C, T>G). In addition to these, CpG>TpG can be included to account for the differential mutation rate of CpG sites ^2^.Hence, we annotated each DNM as one of 7 possible pyrimidine substitutions, counted the occurrence of each substitution per trio, and obtained the proportion of each substitution out of the total DNMs per trio.

Associations were run using generalised linear models (R glm() function) from the quasi Binomial family (logit link), as shown in **Model 4**, and over the same set of non-admixed trios as in the previous section:

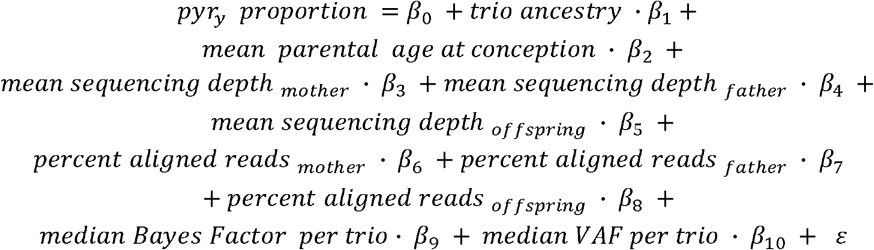

Where *pyr*_*y*_ represents the per trio proportion of DNM being classified in class *y* (i.e.C>A, C>G, C>T, T>A, T>C, T>G, or CpG>TpG). As before, we iteratively changed the ancestry baseline until we had tested all possible ancestry combinations. P-values associated with each ancestry coefficient were corrected to account for multi-testing by pulling together all pyrimidine substitution tests as well (i.e. accounting for 10 ancestry comparisons per each of 7 pyrimidine substitutions, so 70 tests).

### Analysis of smoking and DNM rate

We derived a proxy for the binary “*ever_smoked*” phenotype (0|1) using ICD10 codes available in the secondary care (admitted patient care - APC) data as part of the hospital episode statistics (HES) records available for 100kGP participants ^37^. We identified individuals with at least one ICD10 code related to tobacco smoking behaviour. Specifically, we included Z58.7 (*“Exposure to tobacco smoke”*) and F17-derived codes (*“Mental and behavioural disorders due to use of tobacco”*), with most of the records falling under F17 codes (n F17 codes = 2,165; n Z58 codes = 4). We note that F17 was renamed as “Nicotine dependence” in the 2024 version of ICD10, and that the code Z72, representing “tobacco use”, was not present in the EHRs. We classed individuals having >= 1 smoking ICD10 entry as smokers (i.e. *ever_smoked* = 1) or non-smokers (i.e. *ever_smoked* = 0).

With this, we first built an integrative model of DNM rate including both smoking and ancestry. From the set of non-admixed trios (n trios = 9,833), we further kept trios where at least one parent had APC data available (n trios = 9,031). From these, in 6,221 trios both parents had APC data available, 205 had it for the father only, while 2,605 had it for the mother only. Together with the individual level “*ever smoked”* annotation produced before, for each trio we encoded a 4-level factor called “*parental smoking*”. This indicates whether both parents smoked (n = 293), only the father smoked (n = 664), only the mother smoked (n = 833), or none smoked (n = 7,241). Given that APC data was available for a single parent in 2,810 trios, our “*parental smoking*” encoding assumes that the parent missing APC information is a non-smoker. We re-ran **Model 1** while adding the “*parental smoking*” covariate.

### Refining smoking effect estimate using phased DNM data

From 20,245 non-hypermutator parents with phased DNM information (previously produced by Kaplanis et al. ^4^), and complete metadata we were able to retrieve at least one ICD10 entry for 15,732 individuals (n fathers = 6,599 fathers; n mothers = 9,133). Out of 15,732 individuals with ICD10 code information, 2,169 had at least one ICD10 entry relating to nicotine dependence from ICD10 codes Z58.7 or F17 (n fathers = 996 fathers; n mothers = 1,173).

Combining data from mothers and fathers, we ran associations using a generalised linear model of the quasi-Poisson family (r glm() function), where we controlled for covariates that affect both the calling and phasing of DNMs (**Model 5**).

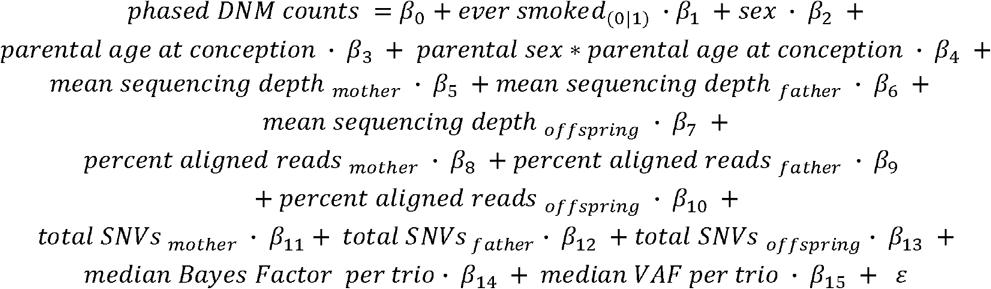

We note that this and other models exclude any effects of cross-parental age on phased mutations (e.g. an effect of maternal age on paternally phased DNMs). This exclusion is justified by the analysis of cross-parental effects presented in **<u>Supplementary Note 5</u>**.

Finally, we ran associations separately for each parent. In such cases we dropped the sex and sex*age covariates while subsetting to either mother or fathers each time. We quantified the effect of smoking in fathers relative to a given change in paternal age using using the formula 4.385+1.296*paternal age, where the intercept and slope were obtained from a simplified negative binomial model keeping all of the covariates in **Model 5** but dropping the smoking effect.

### Derivation of residualised phased DNM counts for genetic associations

We obtained phased autosomal DNM counts across a cohort of 15,885 genetically identified European parents (n fathers = 7,892; n mothers = 7,993; EUR probability >= 0.8) previously produced by Kaplanis et al. ^4^. In addition to the same technical and biological factors accounted for in previous models, we further included the number of SNVs per trio as this would affect the ability of phase DNMs due to the method used for this process ^4^. Hence, as our phenotype, we took the residuals from a generalised linear model from the quasi-Poisson family (**Model 6 - sex combined**):

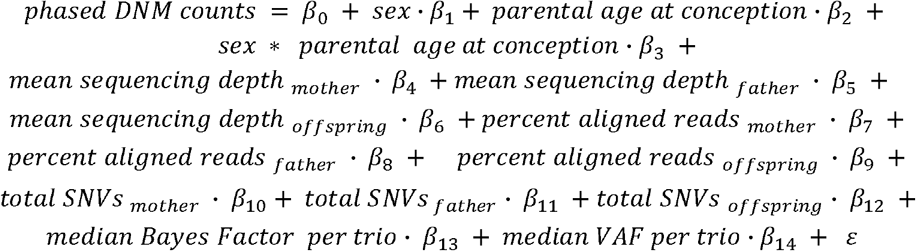

To attempt to identify sex specific differences on DNM rate, we residualized phased DNM counts in a similar way to **Model 6** while subsetting the dataset to either fathers or mothers alone (**Model 7 - sex specific mothers; Model 8 - sex specific fathers**), dropping the sex term and the sex * age at conception interaction term.

Residuals obtained from models 6-8 were later used for heritability estimation and genome-wide association analysis.

### SNP heritability estimation

Genotypes for heritability estimation were processed as follows. Aggregated genotypes from 100kGP project participants had been previously masked to set as missing sites with sequencing depth (DP) < 10, genotype quality (GQ) < 20, or heterozygous genotypes failing the binomial test for allele balance with a p < 10^-3 41^. We restricted the heritability analysis to the 15,885 EUR parents mentioned in the previous section, from which we further removed 478 related individuals (up to 3rd degree) that were previously identified in GEL via the KING genetic relatedness algorithm ^37,42^. From these files, we further removed genotypes with a missing rate > 0.02 and with a Hardy-Weinberg equilibrium test p < 10^−6^.

Heritability was estimated using the residualized phased DNM counts from models 6-8 described in the phased DNM residualisation section (i.e. both parents combined, fathers alone, and mothers alone). Each individual phenotype subset was matched to its respective genotype subset (e.g. fathers-only residualized DNMs to fathers-only genotypes) and 20 population-specific PCs (EUR) were included in the heritability estimation regression for all methods ^37^.

GREML-LDMS was used to calculate heritability across six variance components ^22^. For this, genetic relatedness matrices (GRMs) were calculated using three MAF bins (MAF >= 0.001 & < 0.01, MAF >= 0.01 & < 0.05, and MAF >= 0.05), and two LD bins (low and high LD). LD bins were derived from LD scores calculated directly from the genotype data mentioned above for all variants with MAF >= 0.01 in a 200Kb window. Low LD was defined as those variants having a LD score lower than the genome wide median (median LD score = 81.31), or higher than this in the case of high LD.

### Genome-wide association study for DNM rate

We ran a genome-wide association study for DNM rate on common autosomal SNVs and INDELs (MAF >= 0.05). For this we used the residualized phased DNM counts (models 6-8 in the DNM residualization section) and the linear mixed model implementation of SAIGE v1.0.7 ^23^. Genotype data corresponded to that in the heritability estimation section (including related individuals), with three individual subsets: mothers only (n=7,993), fathers only (n=7,892), and both parents combined (n=15,885). As before, each phenotype subset (residualized phased DNMs) was matched to its corresponding genotype subset.

### Mendelian randomization

To determine causal relationships between different exposures and DNM rate, we ran two-sample Mendelian randomization (MR) analysis using the inverse-variance weighted, MR Egger, and the simple and weighted median-based approaches ^25^. Instrumental variables for the putative exposures were selected from publicly available GWASs. The selected risk factors and their respective sources are shown in **<u>Supplementary Table 3</u>**. These were:

- Age of menopause, selected on the basis of the results in Stankovic et al., 2022^12^, and considered a positive control.
- Smoking initiation, smoking cessation and age of smoking initiation, chosen based on epidemiological associations between smoking and male infertility ^24^ and based on our own results which suggest an association between smoking and DNM rate
- Alcohol use (drinks per week), chosen based on epidemiological associations between alcohol use and male infertility ^24^
- Three phenotypes chosen through a phenome-wide association study of the top SNP from our sex-combined DNM rate GWAS (rs71599241, p-value = 1.01×10^−7^) conducted using an atlas of genetic associations in UK Biobank ^43^. We selected three phenotypes that were nominally significantly associated with this SNP and seemed relevant to reproduction: hydrocele and spermatocele (ICD10 code N43) (p=0.001), diseases of male genital organs (ICD10 codes N40-51) (p=0.004), and sleep duration (p=0.009) ^24^.

As linkage disequilibrium (LD) between instrumental variables can bias MR causal effect estimates due to horizontal pleiotropy ^25^, we pruned the instrumental variables (LD r^2^>0.1). The total number of instrumental variables used per risk factor and LD threshold is shown in **<u>Supplementary Table 3</u>**. The effects of these variants on paternal, maternal, or sex-combined DNM rate were estimated in the GWASs described in the previous section. The full summary statistics for all exposure phenotypes and parameter combinations used for this analysis are included in **<u>Supplementary Table 6</u>** for paternal DNM rate, and **<u>Supplementary Table 7</u>**, for maternal DNM rate, and **<u>Supplementary Table 8</u>** for the sex-combined DNM rate.

## Data availability

Whole-genome sequence data and phenotypic data from the 100,000 Genomes project can be accessed by application to Genomics England (https://www.genomicsengland.cgfbo.uk/research/academic/join-gecip). GWAS summary statistics of DNM rate (sex-combined, maternal only, and paternal only) generated in this study are available as part of our **<u>Supplementary Material</u>**. Publicly available GWAS summary statistics can be accessed at various resources: http://geneatlas.roslin.ed.ac.uk, https://conservancy.umn.edu/handle/11299/241912, https://www.reprogen.org/. Somatic mutations from ascertained smoker individuals can be accessed at: https://data.mendeley.com/datasets/b53h2kwpyy/2. Reference single base substitution mutational signatures used for deconvolution can accessed at: https://cancer.sanger.ac.uk/signatures/sbs/

## Code availability

Plink (v.1.9) was used to process genotype data. Samtools (v.1.15.1) and jvarkit (https://github.com/lindenb/jvarkit) were used to extract read level information. SNP heritability was estimated using GCTA (v1.94.0). SAIGE (v.1.07) was used to perform GWAS. Instrumental variable LD-pruning was performed using LDlinkR (v1.2.2). Two sample Mendelian Randomization was performed using the MendelianRandomization (v.0.9.0) R package. *De novo* mutational signature annotation was performed using the “hdp” R package (https://github.com/nicolaroberts/hdp). Mutational signature deconvolution was performed using SigProfilerExtractor. All remaining analyses were performed in R (v.4.2.1).

## Supplementary Figures

**Supplementary Figure 1.**
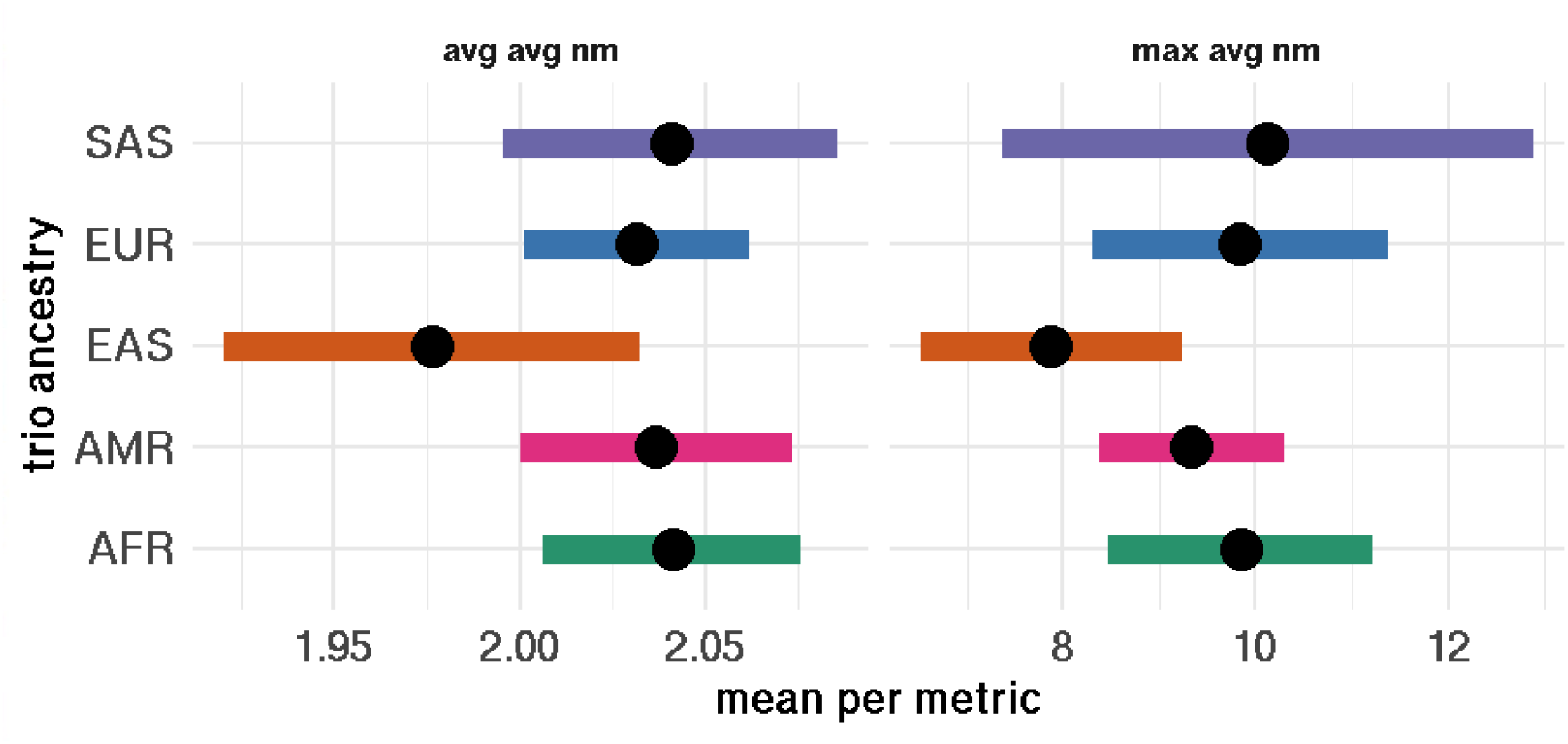
Comparison of means for two metrics summarising the average number of mismatches per read (avg NM) covering the alternate allele at a DNM per trio across ancestries. Left side: Average avgNM per site per trio. Right side: Maximum avgNM per site per trio. Error bars represent 95% confidence intervals for each metric’s mean.

**Supplementary Figure 2.**
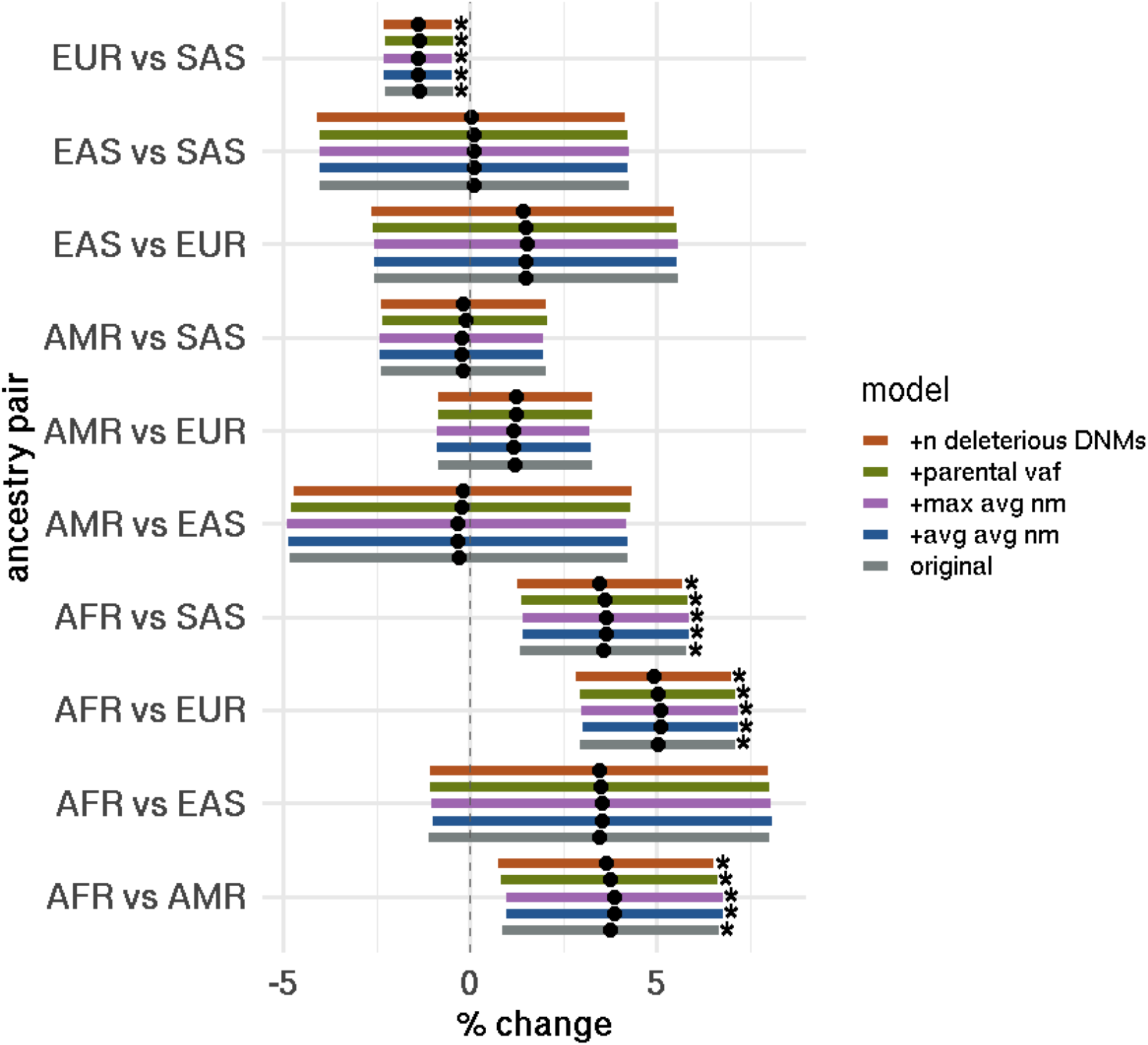
Comparison of number of DNMs between ancestry groups before and after controlling for covariates that aim to capture potential artifacts. Bar colours correspond to a regression model for DNM rate ∼ ancestry, including all of the original covariates plus an extra potential artifactual source. The extra covariates included were average-average mismatches per read per trio (avg avg NM), maximum average mismatches per read per trio (max avg NM), mean parental variant allele fraction per trio (parental VAF), or number of potentially deleterious *de novo* variants (n deleterious DNMs). The grey bar corresponds to the original model outlined in Methods. Asterisks indicate ancestry effect significance at 5% FDR (p adjusted <= 0.05).

**Supplementary Figure 3.**
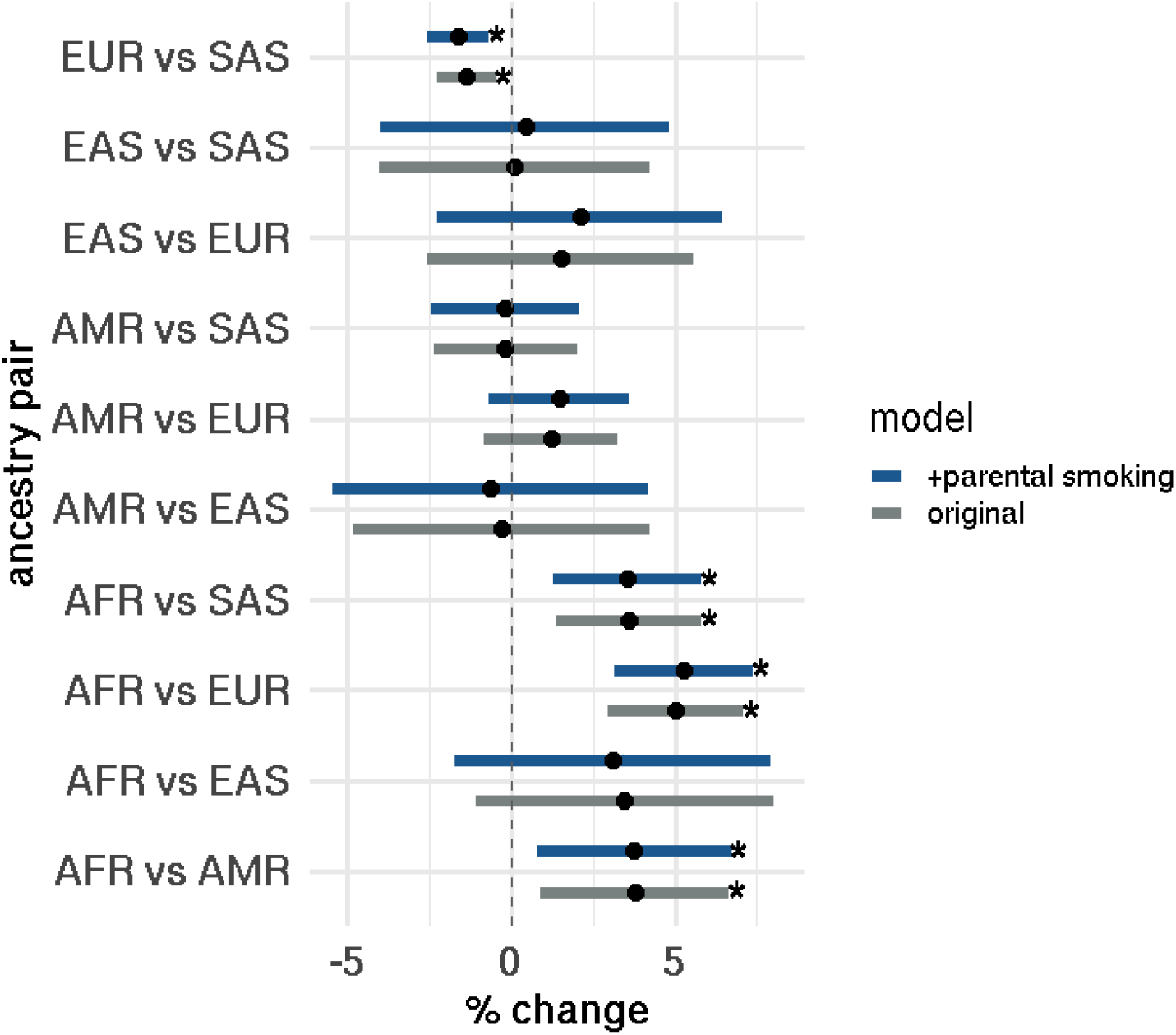
Ancestry effects on DNM rate with and without controlling for parental smoking behaviour. Asterisks indicate significance of the ancestry effect at 5% FDR (p_adjusted_ <= 0.05)

**Supplementary Figure 4.**
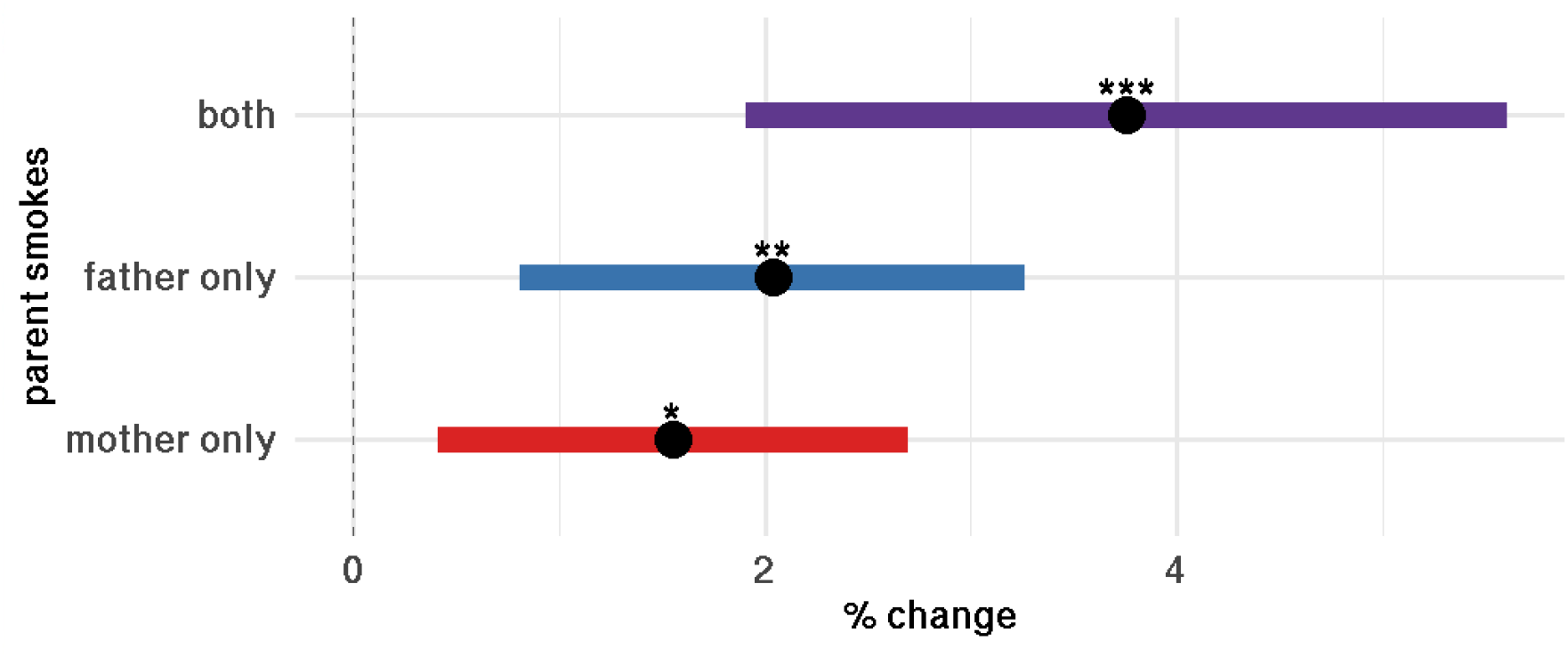
Effect of parental smoking behaviours on DNM rate at the trio level (i.e. using total DNM count). Percentage changes are obtained from the rate ratio effect estimate when comparing smoker parent categories (mother only, father only, both) vs the baseline (no parent smokes). *p<=0.05, **p<=0.005, ***p<=0.0005

**Supplementary Figure 5.**
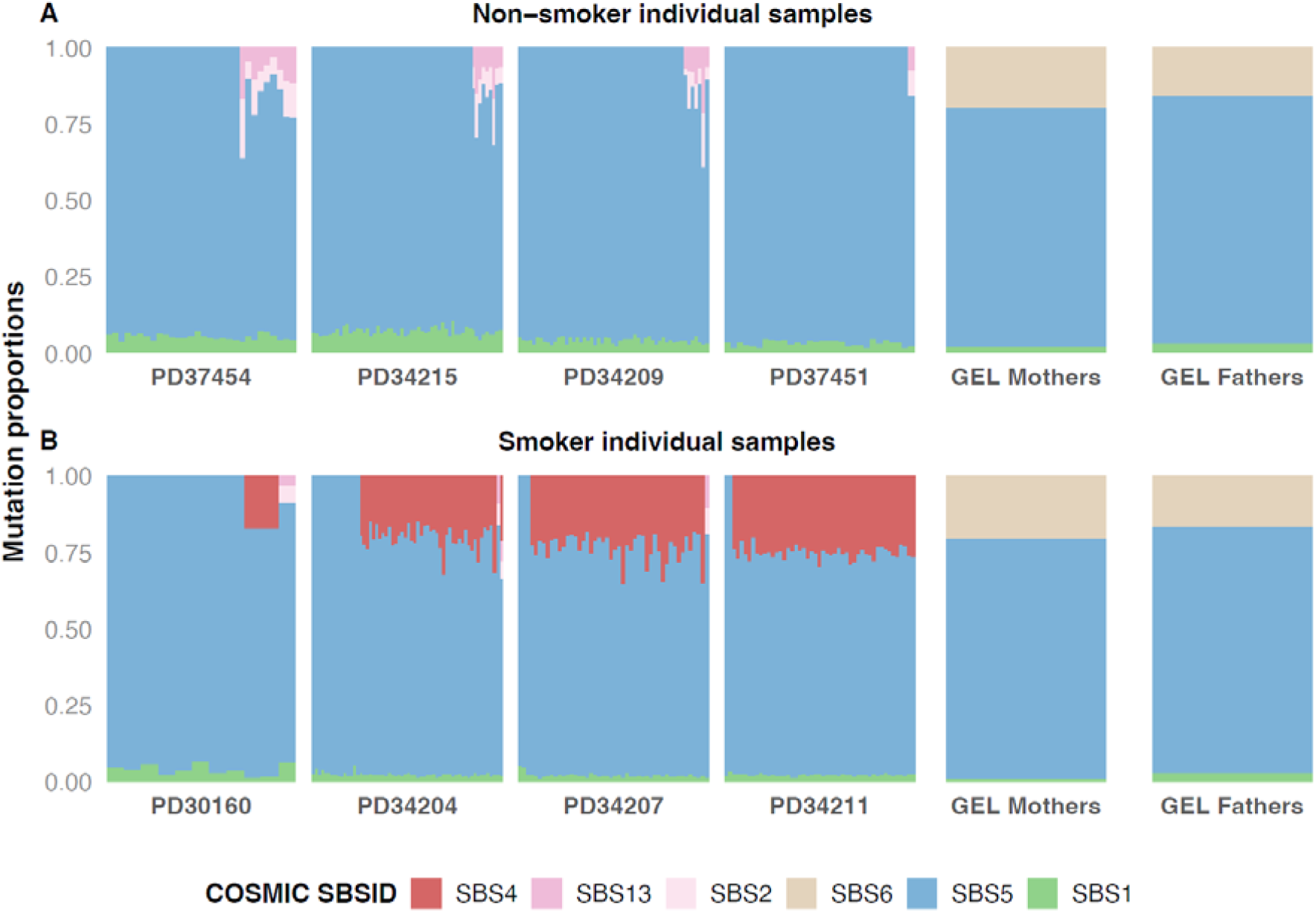
Per-sample mutational signature exposures for A) non-smoker and B) smoker individuals or “meta-individuals” (see **<u>Supplementary Note 4</u>** for methods). Each bar represents the total proportion of mutations assigned to each indicated single base substitution (SBS) signature, amongst somatic mutations detected in bronchial epithelium from four current smokers and four non-smokers from Yoshida et al., 2020 ^44^, or amongst DNMs from “meta-individuals” composed pools of GEL individuals. Mutation exposures in each sample were deconvoluted into six known mutational processes from the COSMIC v3 catalogue ^21^(SBS*N*).

**Supplementary Figure 6.**
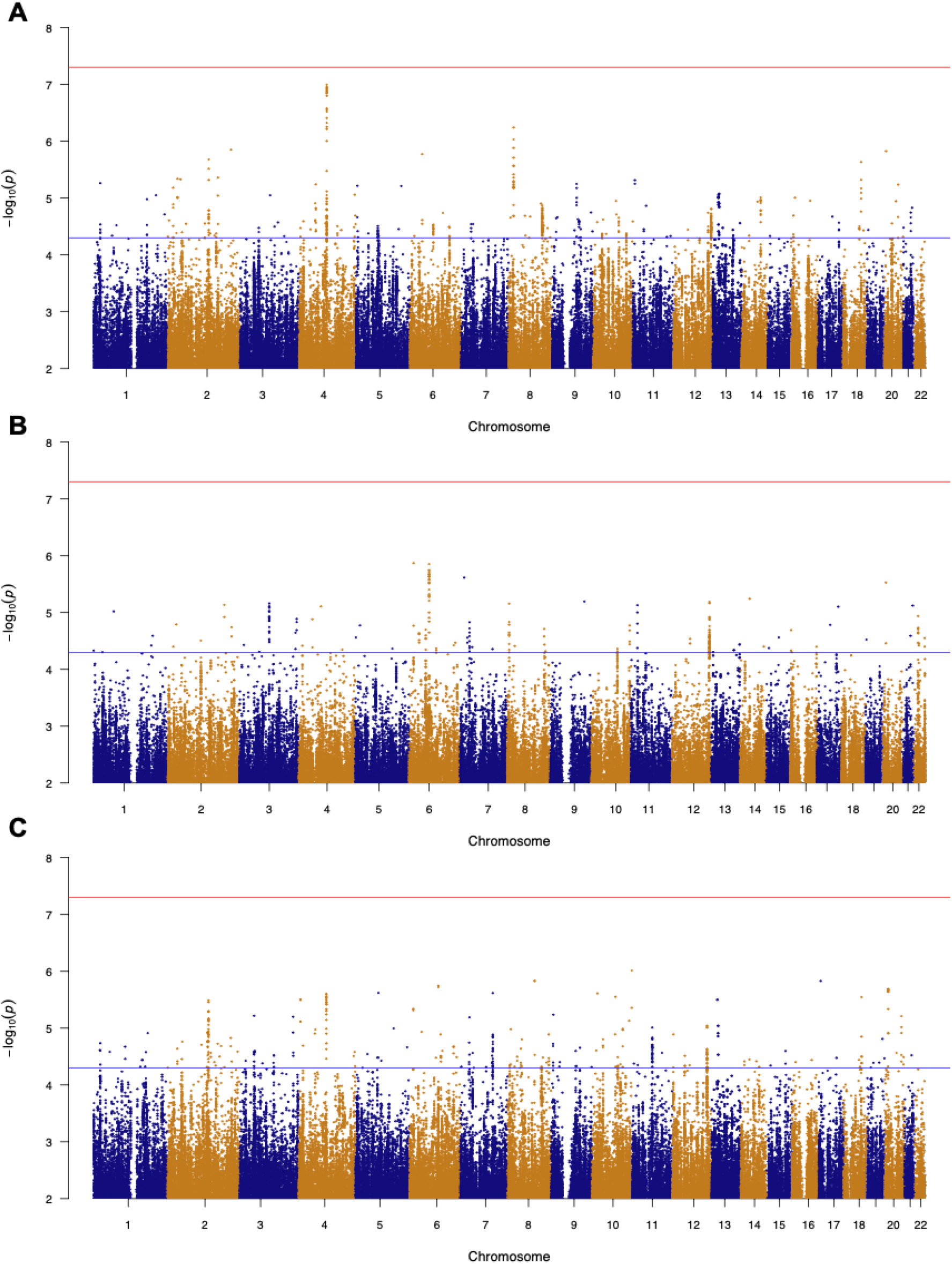
GWAS Manhattan plots for DNM rate in the sex-combined model including both parents (A), fathers only (B), or mothers only (C). Blue line corresponds to suggestive significance (p<=5×10^−5^) while the red line corresponds to the genome-wide significance threshold (p<=5×10^−8^).

**Supplementary Figure 7.**
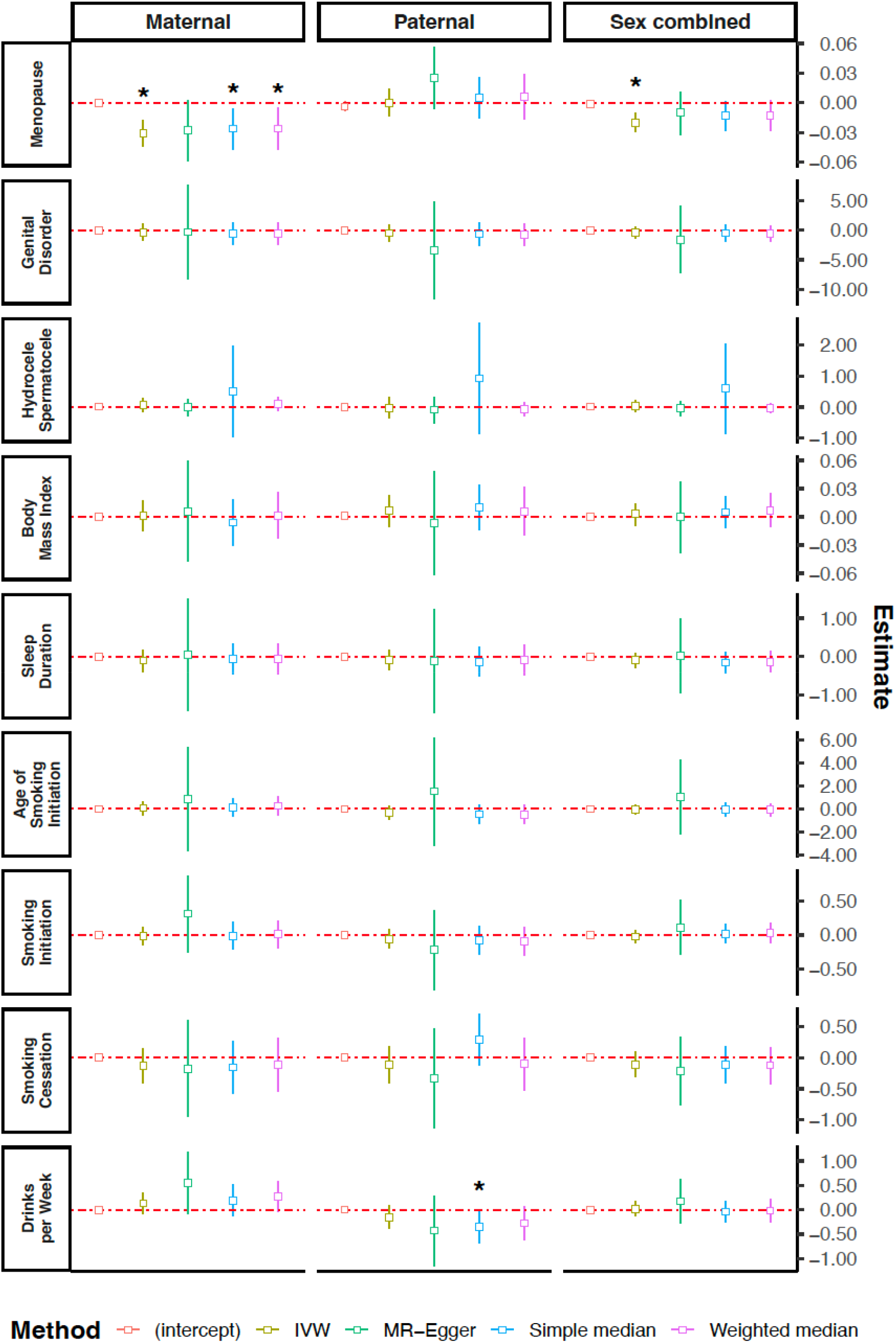
Forest plots showing estimated causal effects of putative risk factors on phased DNM counts obtained with Mendelian Randomisation. This was conducted on maternally phased DNM counts (left), paternally phased DNM counts (centre), and phased DNM counts in both sexes combined (right). Estimates are computed by four different methodologies (colours): IVW, MR-Egger, Simple median and Weighted median. The intercept from MR Egger is also shown (as a test of directional pleiotropy). Bars correspond to 95% confidence intervals for each estimate. Asterisks indicate nominal statistical significance (p <= 0.05). The exposures considered included: age of menopause (“menopause”), diseases of male genital organs (“genital disorder”), sleep duration, hydrocele and spermatocele, smoking initiation (i.e. ever smoked versus never smoked), smoking cessation (i.e. being a current rather than former smoker), age at smoking initiation, drinks per day, and body mass index (BMI). The nominally significant results imply the following directions of effect: later age of menopause causes lower DNM rate in females, and increased number of drinks per week causes a lower DNM rate in males.

**Supplementary Figure 8.**
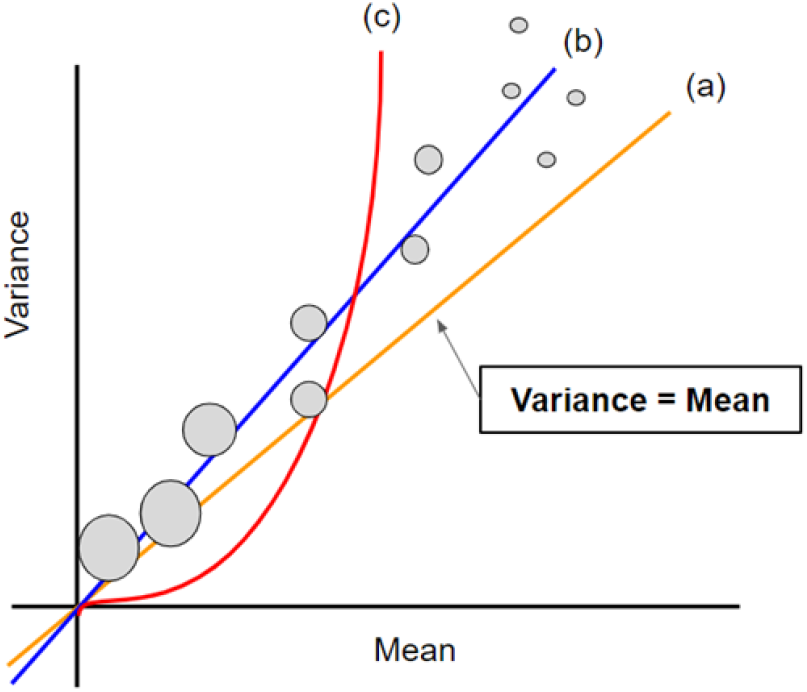
Example of data following various different distributions in the Poisson family. The lines represent (a) Poisson distribution, (b) quasi-Poisson distribution, (c) negative binomial distribution.

**Supplementary Figure 9.**
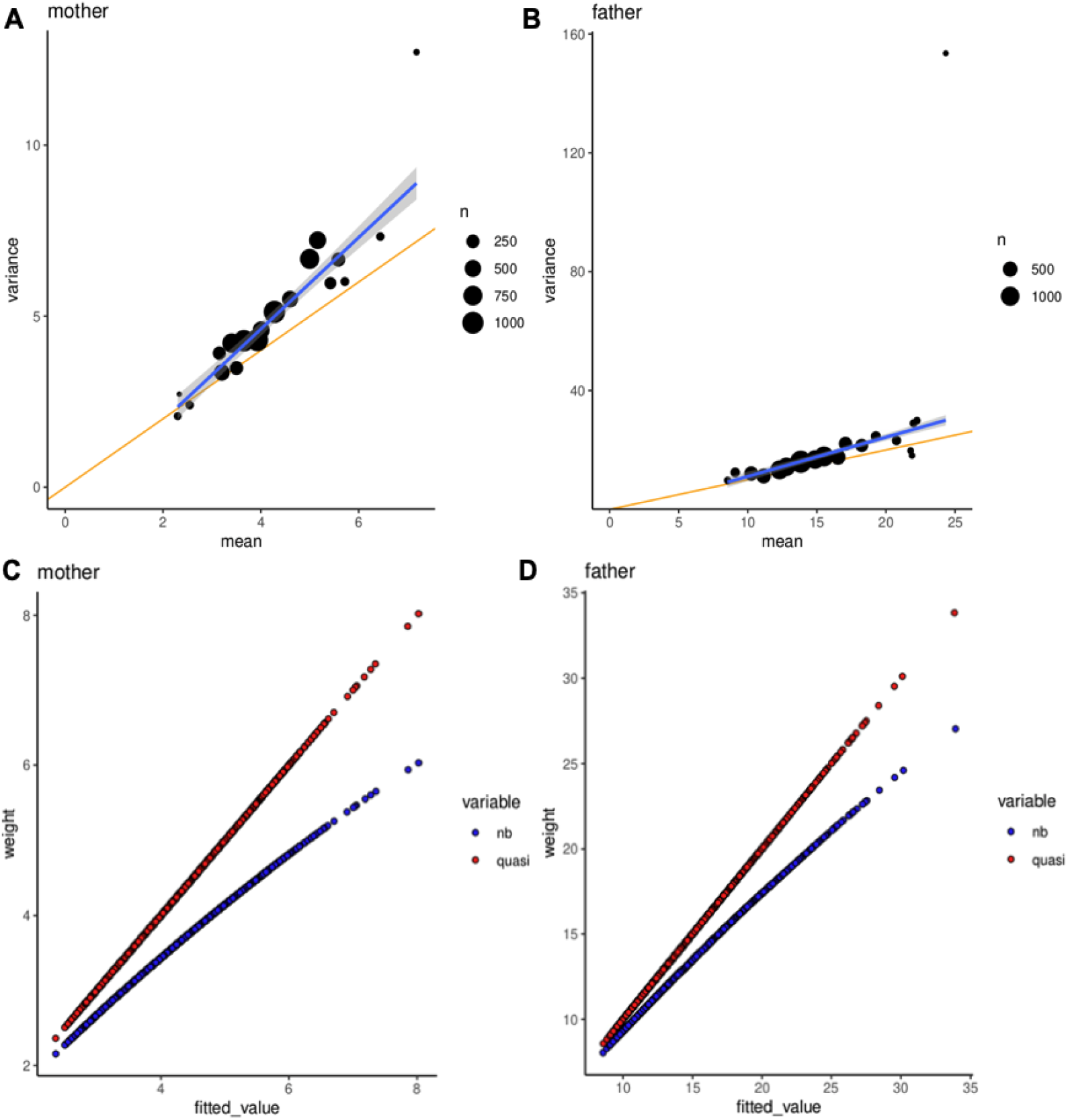
(A-B) Estimated variance-to-mean relationship for phased DNMs in mothers (A) and fathers (B), with a robust regression fit line (blue). The yellow line represents variance = mean line. The size of the circle represents the number of samples within each binned age group. Note that the rightmost points (which are outliers) contain only a tiny number of samples. (C-D) Estimated regression weights as a function of the mean for phased DNMs in mothers (C) and fathers (D).

**Supplementary Figure 10.**
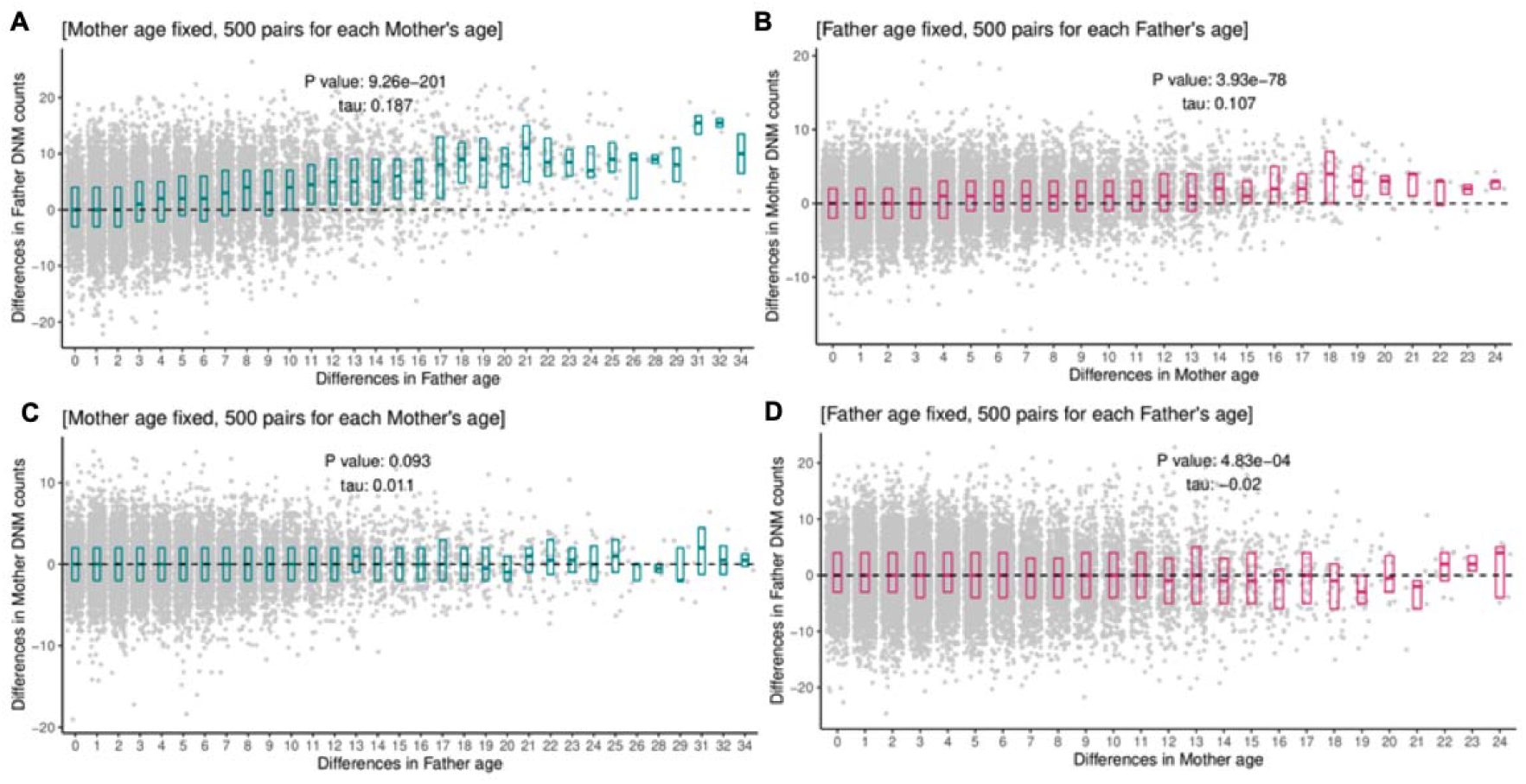
Pairwise comparison of phased DNM counts conditional on the same parental age: each point represents a pair of trios, with 500 pairs per age group in each case. The x-axis displays the difference in paternal ages (a), (c) / maternal ages (b), (d) and the y-axis illustrate the difference in paternal (a), (d) or maternal (b), (c) mutation counts. The boxplots represent 25^th^, 50^th^, and 75^th^ percentile. Tau and p-values are evaluated by using Kendall’s rank correlation test statistic. The correlations observed in panels (a) and (b) correspond to the expected effects of paternal age on paternal DNMs and maternal age on maternal DNMs respectively. Panel (c) corresponds to an effect of paternal age on maternal DNMs and panel (d) corresponds to an effect of maternal age on paternal DNMs.

## Supplementary material

### DNM rate GWAS summary statistics

All these files include the columns: *“chrom”*: chromosome, *“pos”*: GRCh38 assembly position. *“marker*.*id”*: Construct of “chrom:pos_other.allele_effect.allele”. *“other*.*allele”*: allele NOT used for regression estimate. “*effect*.*allele”*: Allele used as reference for beta effect; “*af_effect*.*allele*”: Sample subset specific allele frequency of the effect allele. “*Beta*”: Regression beta effect (SAIGE); “*se*”: Standard error for beta effect; “*p*.*value*” P value for beta effect.

- **Sex-combined DNM rate GWAS:** DNM rate GWAS summary statistics for the sex combined individual set.
- **Paternal DNM rate GWAS:** paternal DNM rate GWAS summary statistics for the fathers subset.
- **Maternal DNM rate GWAS:** maternal DNM rate GWAS summary statistics for the mothers subset.

## Supplementary Tables

ancestral_gen_and_env_fctrs_dnmrate_supplementary_tables.xlsx

**Supplementary Table 1**

**Summary statistics from testing for ancestry differences in DNM counts using generalised linear models**. We conducted pairwise comparisons across five super-continental ancestry classifications (AFR, AMR, EAS, EUR, SAS) using quasi-poisson models. *“reference ancestry”*: ancestry used as baseline for given comparison; *“compared ancestry”*: ancestry compared against baseline; *“estimate”*: generalised linear model estimate (in log(rate ratio) scale) corresponding to the effect of being a member of the “*compared ancestry*” rather than the “*reference ancestry*” group; “*se*”: standard error for the estimate effect; “*nominal p*”: p value for the effect estimate; “*fdr padj”*: nominal p for all ancestry comparisons adjusted by the “Benjamini–Hochberg” method (FDR); “*significance label*”: asterisks indicate comparisons with an “*fdr padj*” <= 0.05 (FDR 5%).

**Supplementary Table 2**

**Summary statistics from testing for ancestry differences in DNM spectra (pyrimidine substitution counts/total trio DNMs) using generalised linear models**. We conducted pairwise comparisons across five super-continental ancestry classifications (AFR, AMR, EAS, EUR, SAS) using quasi-Binomial models. *“reference ancestry”*: ancestry used as baseline for given comparison; *“compared ancestry”*: ancestry compared against baseline; “*pyr subs*”: pyrimidine substitution proportion being compared; *“estimate”*: generalised linear model estimate (in log(rate ratio) scale) corresponding to the effect of being a member of the “*compared ancestry*” when comparing against the “*reference ancestry*” group; “*se*”: standard error for the estimate effect; “*nominal p*”: p value for the effect estimate; “*fdr padj”*: nominal p for all ancestry comparisons adjusted by the “Benjamini–Hochberg” method (FDR); “*significance label*”: asterisks indicate comparisons with an “*fdr padj*” <= 0.05 (FDR 5%).

**Supplementary Table 3**

**Number of SNPs used as instrumental variables (IVs) in Mendelian Randomisation (MR) analyses**. “*Phenotype*”: Exposure phenotype; *“Study Origin”*: Publication of origin of exposure IVs. “*Sex-combined*”: number of IVs extracted from the sex-combined DNM rate GWAS; “*Father*”: number of IVs extracted from the paternal DNM rate GWAS; “*Mother*”: number of IVs extracted from the maternal DNM rate GWAS.

**Supplementary Table 4**

**Summary of number of mutations and samples used for signature extraction and deconvolution to detect potential mutational signatures associated with smoking**. “*Origin*”: Sample origin, either GEL (pooled individuals in this study), or Yoshida et al. 2020 for our external reference sample set; “*donor*”: Donor name for each of the sample sets shown in Supplementary **Figure 4** (X-axis identifier); “*smoker status*”: smoker status of the sample in question, for samples taken from Yoshida et al. 2020^44^ samples, “non-smoker” corresponds to the “never smoker” tag in the original publication, while “smoker” corresponds to the “current smoker” tag, respectively; “*n inds pooled*”: total individuals pooled for each of the GEL meta-samples; “n samples” number of samples included for each donor (bronchial epithelium samples in the case of Yoshida et al., 2020 samples), set to NA for GEL meta-individuals; “*n mutations*”: Total number of mutations included for each donor, in the case of GEL samples, this corresponds to the number of pooled DNMs per meta-individual.

**Supplementary Table 5**

**Pyrimidine substitution differences across continental populations for SNP data from Harris and Pritchard, 2017** ^14^. *“reference ancestry”*: ancestry used as baseline for given comparison; *“compared ancestry”*: ancestry compared against baseline; “*pyr subs*”: pyrimidine substitution proportion being compared; “*OR”:* proportion differences obtained directly from counts collapsed into N-pyrimidine substitutions; “*pval*”: p values from chi-squared test comparison of pyrimidine substitution counts between two populations; “*ordered p*”: chi-squared test p values calculated for pyrimidine substitutions arranged by raw significance; “*significance label*”: comparisons with ordered p value <= 1e-4; “pyr code”: number of pyrimidine substitutions used to collapse the original 96 substitution code annotation. This can be either a 7-code (C>A, C>G, C>T, CpG>TpG, T>A, T>G, and T>C; note that C>T includes CpG>TpG) or a 6-code excluding the CpG>TpG class (i.e. not making distinction of C>T substitutions occurring in CpG sites vs those occurring elsewhere).

**Supplementary Table 6**

**Mendelian Randomisation summary statistics for paternal DNM rate. “***Method”:* Method used for Mendelian Randomisation. *“Std*.*Error”:* Standard error for MR effect estimate. *“95% CI lower”:* 95% confidence interval for MR effect estimate. *“95% CI upper”:* 95% confidence interval for MR effect estimate. *“P-value”:* MR effect estimates p value. *“Exposure”: Tested phenotype exposure. “LD”:* Linkage disequilibrium threshold for instrumental variable pruning.

**Supplementary Table 7**

**Mendelian Randomisation summary statistics for maternal DNM rate**.

Column contents match those of Supplementary Table 8.

**Supplementary Table 8**

**Mendelian Randomisation summary statistics for sex combined DNM rate**.

Column contents match those of Supplementary Table 8 and 9.

## Supplementary Notes

### Supplementary Note 1

#### Modelling mean-variance overdispersion for DNM count data

Given that the phenotype of interest (DNM rate) in our study represents count data, a natural assumption would be that it follows a discrete distribution within the Poisson family. Overdispersion can be modelled by different distributions such as Poisson, Quasi-poisson, and negative binomial, as shown in **Supplementary Figure 8**. The quasi-Poisson distribution (line b) assumes the variance is a linear function of the mean, same as the Poisson distribution (line a) but with a slope greater than 1, and the negative binomial distribution posits a quadratic relationship between the mean and variance (line c).

To elucidate the form of overdispersion present in the phased DNM count data, we constructed a diagnostic plot of the empirical fit of the variance-mean relationship, grouping samples by parental age at conception (the major factor determining DNM rate) into twenty bins of equal age intervals. We calculated the mean of samples falling into each bin and placed them on the x-axis, while the sample variances were computed and placed on the y-axis. This plot can be observed in **Supplementary Figure 9** (A,B). Here, it can be noted that the data exhibits a linear relationship with a steeper slope than the variance = mean line. This linear trend suggests that the quasi-Poisson distribution is a more suitable model for our data than the negative binomial or Poisson distributions.

Additionally, to fit the model to the data, both quasi-Poisson and negative binomial models employ the iteratively weighted least-squares algorithm. The concept of weight here was used to take into account the unequal variance among residuals by modelling the objective function in a form of weighted least squares problem. Thus, another distinguishing factor between the quasi-Poisson and negative binomial regression models is the difference in the weights between these models. For instance, in the quasi-Poisson model, the weights are directly proportional to the mean, whereas in the negative binomial model, the weights exhibit a concave relationship with the mean, as shown in the equation below.

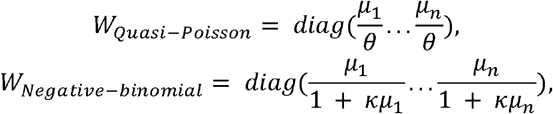

We performed both quasi-Poisson and negative binomial regression analyses on the data sets for paternally and maternally phased DNMs, respectively. In each model, we plotted the estimated weights against the fitted values. As seen in **Supplementary Figure 9** (C,D), the quasi-Poisson regression results exhibit a linear relationship, characteristic of the quasi-Poisson model, while in the case of the negative binomial model, a slight curvature is observed. Given the patterns observed in the relationship between variance and mean, as well as the changing patterns of weights, we conclude that the quasi-Poisson model is more suitable for our data ^45^.

### Supplementary Note 2

#### Ensuring that the associations between ancestry and DNM rate are not due to technical artefacts or ascertainment bias

We wanted to rule out that observed differences in DNM counts between ancestries were due to differences in the mapping quality of the reads resulting from ancestral biases in the reference genome. To test this, we first calculated the average number of mismatches of reads containing the alternate allele per DNM (avgNM), then calculated the mean and maximum avgNM per offspring. We fit an ordinary least squares model to compare the mean and maximum avgNM values per trio between the different ancestry classes, as follows:

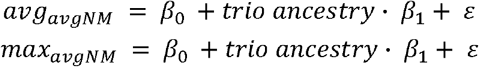

This analysis found that ancestry was not associated with either the average of the maximum avgNM per offspring (**Supplementary Figure 1**). We separately included the mean and maximum avgNM values per trio as extra covariates in **Model 1** (main Methods) section to check whether this significantly altered the original estimates, which it did not (**Supplementary Figure 2**).

Another indicator of potential mapping bias may be the parental coverage for the alternate site classed as “*de novo*’’ in the offspring. If a given ancestry group has higher mapping errors on average, variants which are present in the parents and passed on to the child may have low variant allele fraction (VAF), such that these sites are erroneously called as DNMs. To check this, we calculated the mean parental variant allele fraction (VAF) ([maternal VAF + paternal VAF] / 2) at putative DNM sites, calculated the mean across DNMs per offspring, and then included this metric as a covariate in **Model 1 (**main Methods). We found that this does not affect the original ancestry effect estimates (**Supplementary Figure 2**).

Finally, it is conceivable that there may have been ascertainment biases during recruitment to the 100,000 Genomes Project such that families from certain ancestry groups were more likely to be recruited if their affected child had a pathogenic DNM rather than some other genetic or non-genetic cause. This would be expected to manifest in differences in the number of deleterious DNMs (most of which are probably protein-altering) between ancestries, which could, in theory, drive the ancestral differences we see in overall DNM rate. We counted the number of protein-coding DNMs per proband (including those with worse consequence *“missense_*”, “start_lost”, “stop_lost”, “stop_gained”, “stop_retained”*, or *“splice_*”*) and included this as an extra covariate in **Model 1** (main **<u>Methods</u>**). The observed ancestry associations to DNM rate remained unchanged, suggesting that these were not due to ancestry-related ascertainment biases (**Supplementary Figure 2**).

### Supplementary Note 3

#### Comparing ancestry-associated differences on mutation spectra using DNM data and polymorphism data

Ancestry-associated differences in germline mutation spectra were previously reported by Harris and Pritchard ^14^. These were discovered using common polymorphism data from the 1000 Genomes Project. Briefly, each SNP in that dataset was classified according to its ancestral and derived allele (C>A, C>T, C>G, A>T, A>G, A>C), and the base pair immediately 5’ and 3’ of it ^14^. This results in each SNP being classified into 1 of 96 possible combinations of flanking context and ancestral-to-derived allele substitutions. Counts for each of the 96-substitutions were generated for each of the five continental super-populations in the 1000 Genomes Project (AFR, AMR, EAS, EUR, SAS). Ancestry-associated differences in spectra were calculated as follows. First, for a given substitution (e.g. “A[C>T]A”), population-specific proportions were calculated as the ratio between the substitution of interest over the sum of the counts for the rest of the substitutions in that population. Then, for each substitution category, population-specific proportions in two populations of interest (e.g. EUR and SAS) were compared.

We compared our mutational spectra results with the findings described by Harris and Pritchard. For this, we obtained the supplementary data corresponding to **Supplementary Figure 1** of their paper ^14^, which consisted of a 5×96 count matrix (ancestry x substitution category). We changed the encoding of the 96-substitution code from this data to make it compatible with our own annotations. Due to limited power in our study, we were not able to consider the full 96-substitution code, so we collapsed counts in the 96-substitution code presented in their paper into a 6-pyrimidine substitution code (switching the substitution to the complementary strand where necessary so we considered only C>T, C>A, C>G, T>A, T>G, and T>C variants). Finally, to obtain p-values for each ancestry comparison and pyrimidine substitution, we followed the same method described by Harris and Pritchard. Briefly, we first obtained chi-square p-values for all population-pairs and substitution comparisons using 2*2 count matrices as shown below:

|  |  |
| --- | --- |
| $S_{p1}^{(m)}$ | $T_{p1} - S_{p1}^{(m)}$ |
| $S_{p2}^{(m)}$ | $T_{p2} - S_{p2}^{(m)}$ |

where *S*_*p*l_^(*m*)^ represents the number of substitutions of type *m* in population *x*, and *T*_*px*_ represents the total number of substitutions in that population.

Then, for each population pair, we obtained ordered p-values by iteratively comparing counts for the substitution with lowest p-value against the counts of the substitution with the next lowest p-value as illustrated below.

|  |  |
| --- | --- |
| $S_1^{(m_i)}$ | $\sum^{n_{pyr\ subs}}_{j=i+1} S_1^{(m_j)}$ |
| $S_2^{(m_i)}$ | $\sum^{n_{pyr\ subs}}_{j=i+1} S_2^{(m_j)}$ |

where *m*_*i*_ represents the substitution with the lowest p value, *m*_*j*_ represents the substitution for the next lowest p-value, and *n*_*pyr subs*_ represents the size of the substitution code being tested (e.g 6 pyrimidine substitutions). As in Harris’ work, all comparisons with an ordered p-value <= 1×10^−5^ were deemed to be significant.

We compared the results from Harris and Pritchard against the significant spectrum changes that we identified in our study (main **Figure 1B**), specifically for SAS and EUR pairs. Their data showed a significant enrichment of C>A (OR = 1.011, ordered p = 2.22×10^−20^), and a depletion of T>C (OR = 0.994, ordered p = 1.10×10^−14^) and T>G (OR = 0.987, ordered p = 1.05×10^−25^) substitutions (**Supplementary Table 5**). Aiming to account for the differential mutation rate produced by the spontaneous cytosine deamination occurring in CpG islands ^19^, we defined a category CpG>TpG for C>T sites occurring next to a CpG site. We found that CpG>TpG proportions were enriched in SAS compared to EUR (OR = 1.010, ordered p =3.27×10^−7^; **Supplementary Table 5**). We note that the effect directions in these comparisons are inverted compared to ours since we used SAS ancestry as the baseline. Hence, the differences reported by Harris and Pritchard would be equal to the inverse of the odds ratio they obtained using the EUR ancestry as the baseline. Taking this into account, our study reproduced the depletion of C>A and the enrichment of T>C in EUR relative to SAS.

### Supplementary Note 4

#### Attempting to identify associations between parental smoking behaviour and DNM mutation spectra

We first tested if smoking is a significant predictor of pyrimidine substitution proportion differences in parentally phased DNMs. For this, in the same way to other analyses, we annotated parentally phased DNMs according to their pyrimidine substitution type (i.e. C>A, C>G, CpG>TpG, C>T, T>A, T>C, T>G) and calculated the proportion of DNMs in each substitution group per individual (out of total phased DNMs per individual). Then, we regressed each pyrimidine proportion on the smoking status of the individual, parental age at conception, and different quality control covariates, as shown in the next model:

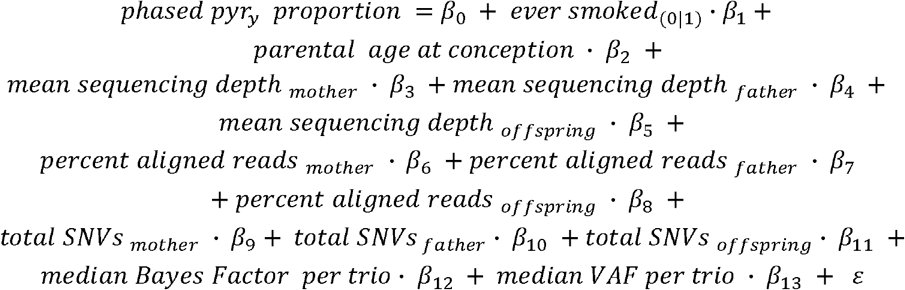

where “*ever smoked*” was derived in the same way as described in the main methods. We restricted this analysis to individuals with DNM phase and EHR data available (n fathers = 6,599 fathers; n mothers = 9,133). We did not find any significant association between being a smoker and any of the tested pyrimidine proportions (all nominal p > 0.05, data not shown).

Next, we asked if known mutational signatures associated with smoking behaviour could be deconvoluted from phased DNMs obtained from parents who smoke. Different somatic mutational processes operating across tissues display distinctive mutational patterns that can be deconvoluted using dimensionality reduction and classification algorithms ^46^. Broadly, this process consists of the steps: 1) generation of mutation counts matrices per sample, with mutations typically annotated using a 96-pyrimidine substitution code ^40,47^, 2) *de novo* signature extraction, which aims to identify single base substitution (SBS) patterns in the input data in an unbiased manner ^40^, and 3) signature decomposition, which aims to compare the identified *de novo* patterns to publicly available annotated SBS signatures, such as the COSMIC database ^21,40^.

To apply these methods to our own data, we first annotated all parentally phased DNMs according to 1) their pyrimidine substitution direction, and 2) their flanking 5’ and 3’ base pair, after which each DNM was classified in 1 of 96 possible SBSs. As mutational signature extraction is a process usually applied to somatic samples (with a mutation load several orders of magnitude higher than that of the germline ^3^), we reasoned that the average number of phased DNMs in a single individual (∼15 for fathers, ∼4 for mothers) may be insufficient to accurately identify mutational signatures. For this reason, we pooled together phased DNMs by smoking status group and sex to create four “meta-individuals”, each representing all of: 1) smoker fathers, 2) smoker mothers, 3) non-smoker fathers, and 4) non-smoker mothers. Then, for each of these synthetic samples, we counted the occurrence of each 96 SBSs. Ninety-nine DNM sites were shared by more than one individual in a given pool, and we removed duplicate variants from each meta-individual before counting. To prevent potential ancestry-related noise, only EUR individuals were considered for DNM pooling. The total number of DNMs and individuals represented by each meta-individual is shown in **Supplementary Table 4**. Aiming to increase our power to pick up any smoking signature signal, we merged our count matrix with counts obtained from an external reference panel containing 337 normal lung (noncancerous bronchial epithelium) samples from four ascertained smokers and four non-smokers ^44^. This matrix was used to identify *de novo* mutational signatures using HDP ^48,49^. The identified *de novo* signatures were compared against the COSMIC SBS v3 database ^21,47^ using the cosine similarity metric implemented in the “lsa” R package ^50^. We kept all *de novo* signatures / COSMIC SBS pairs with cosine similarities >= 0.8, or containing any of the COSMIC signatures reported by Yoshida et al., 2020 ^44^ for the included external samples. With this, we deconvoluted *de novo* signatures into 6 COSMIC SBS signatures using the “*decompose*.*fit*” function implemented in the SigProfilerExtractor software ^40^. Although we were able to correctly identify the hallmark tobacco smoking SBS signature (COSMIC’s SBS4) in the external smoker individuals, we did not identify this in any of the meta-individuals from GEL (**Supplementary Figure 5**).

There could be several reasons behind the apparent absence of any smoking-associated signatures in the GEL smokers. First, we may just be underpowered to detect this since the number of DNMs in our smoker “meta-individuals” is, on average, more than an order of magnitude lower than the average number of somatic mutations in the lung tissue from smokers sequenced in Yoshida et al. (**Supplementary Table 4**). Second, even though it is well established that tobacco smoke is causal for specific mutational signatures in lung tissue ^44^, its effect may not be the same across other tissues, including the germline. On this note, even lung tissue in regular smokers displays some degree of heterogeneity in terms of mutation burden and overall tobacco smoke signature load ^44^. Such heterogeneity may also be expected from tissues not directly exposed to tobacco smoke, potentially even to a higher degree. As it has been shown that tobacco smoke signatures are present in cancerous tissue outside the lung, such as bladder cancers ^51^, we cannot rule out that smoking-associated signatures can be found outside of the lung (even in the germline), but a larger sample size may be required to detect this signal.

### Supplementary Note 5

#### Cross-parental effects on early embryonic mutations

It has been suggested that maternal ageing may lead to an increased frequency of early post-zygotic mutations during the initial cell divisions of the embryo ^52^. Some of these mutations may then appear in some or all offspring somatic cells and be called DNMs. In particular, those occurring on paternally derived chromosomes would be interpreted as paternal DNMs, meaning that such an effect could manifest as an effect of maternal age on the number of paternally DNMs. Gao et al.^52^ found just such a signal in a dataset of 1,548 Icelandic trios ^53^. Replicating their analysis, we investigated within- and cross-parental effects in our much larger cohort, as shown in **Supplementary Figure 10**.

Panel (a) shows the effect of paternal age on paternal mutations, controlling for maternal age. Each point represents a pair of trios in which the maternal age is the same, and where the difference in paternal ages between the trios represented by the x-coordinate and the corresponding difference in phased paternal DNM counts is represented by the y-coordinate. For each maternal age in the dataset, 500 trio pairs were selected at random. As expected, this plot shows an increased differential paternal DNM count with increasing difference in paternal age, corresponding to the well-established effect of paternal age on paternal DNMs. Panel (b) shows the equivalent within-parental effect of maternal age on maternal DNMs, here plotting trio pairs with the same paternal age, for all paternal ages in the dataset.

Panel (c) shows the effect of paternal age on maternal mutations. Here as in panel (a), each point represents a pair of trios in which the maternal age is the same and the difference in paternal ages is represented by the x-coordinate, but now the y-coordinate gives the corresponding difference in phased maternal DNM counts. As expected, no effect of paternal age on maternal DNM count is apparent in this plot. Finally, panel (d) shows the effect of maternal age on paternal DNMs. In the analysis of Gao et al.^52^, this showed a weakly significant positive correlation, but in our larger dataset there is no such signal - in fact there is an apparent negative correlation, caused by the sparsity of data points at large maternal age differences.

Thus these findings are consistent with a limited impact of parental age on the number of early postzygotic mutations in the embryo.

